# Monkey Plays Pac-Man with Compositional Strategies and Hierarchical Decision-making

**DOI:** 10.1101/2021.10.02.462713

**Authors:** Qianli Yang, Zhongqiao Lin, Wenyi Zhang, Jianshu Li, Xiyuan Chen, Jiaqi Zhang, Tianming Yang

## Abstract

Humans can often handle daunting tasks with ease by developing a set of strategies to reduce decision making into simpler problems. The ability to use heuristic strategies demands an advanced level of intelligence and has not been demonstrated in animals. Here, we trained macaque monkeys to play the classic video game Pac-Man. The monkeys’ decision-making may be described with a strategy-based hierarchical decision-making model with over 90% accuracy. The model reveals that the monkeys adopted the take-the-best heuristic by using one dominating strategy for their decision-making at a time and formed compound strategies by assembling the basis strategies to handle particular game situations. With the model, the computationally complex but fully quantifiable Pac-Man behavior paradigm provides a new approach to understanding animals’ advanced cognition.

**One-Sentence Summary:** Macaque monkeys play Pac-Man with strategy-based hierarchical decision making, a cognitive capacity hitherto unknown in them.

## Introduction

Our lives are full of ambitious goals to achieve. Often, the goals we set out to accomplish are complex. For it to be acquiring a life with financial stability or winning the heart of your love of life, these ambitious goals are often beyond the reach of any straightforward decision-making tactics. They, however, may be approached with a specific and elaborate set of basis strategies. With each strategy, individuals may prioritize their gains and risks according to the current situation and solve the decision making within a smaller scope. As we live in a dynamic world that presents us with unexpected disturbances, it is also crucial to have the flexibility to alter our course of strategies accordingly. Additionally, the basis strategies can be pieced together and combined into compound strategies to reach grander goals.

For animals living in nature, the ability to flexibly formulate strategies for a complex goal is equally, if not more, crucial in their lives. Many have shown that animals exhibit complex strategy-like behaviors (Beran, 2015; Bird & Emery, 2009; Brotcorne et al., 2017; Gruber et al., 2019; Leca et al., 2021; Loukola et al., 2017; Reinhold et al., 2019; Sabbatini et al., 2014; Sanz et al., 2010), but quantitative studies are lacking. Moreover, despite the continuing effort in studying complex behavior in animals and the underlying neural mechanisms (Haroush & Williams, 2015; Kira et al., 2015; Ong et al., 2020; Yoo et al., 2020), the level of complexity of the existing animal behavioral paradigms are insufficient for studying how animals manage strategies to simplify a sophisticated task. A sufficiently complex behavior task should allow the animal to approach an overall objective with a variety of strategies in which both the objective, its associated rewards and cost, and the behaviors can be measured and quantified. Establishing such a behavior paradigm would not only help us to understand advanced cognitive functions in animals but also lay the foundation for a thorough investigation of the underlying neural mechanism.

Here, we adapted the popular arcade game Pac-Man. The game was tweaked slightly for the macaque monkeys. Just as in the original game, the monkeys learned to use a joystick to control the movement of Pac-Man to collect all the pellets inside an enclosed maze while avoiding ghosts. The monkeys received fruit juice as a reward instead of earning points. The animals were able to learn how each element of the game led to different reward outcomes and made continuous decisions accordingly. While the game is highly dynamic and complex, the game is essentially a foraging task, which may be the key to successful training. More importantly, both the game states and the monkeys’ behavior were well-defined and could be measured and recorded, providing us opportunities for quantitative analyses and modeling.

The game has a clear objective, but an optimal solution is computationally difficult. However, a set of intuitive strategies would allow players to achieve reasonable performance. To find out whether the monkeys’ behavior can be decomposed into a set of strategies, we fit their gameplay with a dynamic compositional strategy model, which is inspired by recent advances in the artificial intelligence field in developing AI algorithms that solve the game with a multi-agent approach (Foderaro et al., 2017; Rohlfshagen et al., 2017; Sutton et al., 1999; Van Seijen et al., 2017). The model consists of a set of simple strategies, each considers a specific aspect of the game to form decisions on how to move Pac-Man. By fitting the model to the behavior of the animals, we were able to deduce the strategy weights. The model was able to achieve over 90% accuracy for explaining the decision-making of the monkeys. More importantly, the strategy weights revealed that the monkeys adopted a take-the-best heuristic by using a dominant strategy and only focusing on a subset of game aspects at a time. In addition, the monkeys were able to use the strategies as building blocks to form compound strategies to handle particular game situations. Our results demonstrated that animals are capable of managing a set of compositional strategies and employing hierarchical decision making to solve a complex task.

## Results

### The Pac-Man Game

We trained two monkeys to play an adapted Pac-Man (Namco®) game (Figure 1A). In the game, the monkeys navigated a character known as Pac-Man in a maze and their objective is to traverse through the maze to eat all the pellets and energizers. The game presented the obstacles of having two ghosts named Blinky and Clyde, who behaved as predators. As in the original game, each ghost followed a deterministic algorithm based on the Pac-Man’s location and their locations with Blinky chasing Pac-Man more aggressively. If Pac-Man was caught, the monkeys would receive a time-out penalty. Afterwards, both Pac-Man and the ghosts were reset to their starting locations, and the monkeys could continue to clear the maze. If the Pac-Man ate an energizer, a special kind of pellet, the ghosts would be cast into a temporary scared mode. The Pac-Man could eat the scared ghosts to gain extra rewards. All the game elements that yield points in the original game provided monkeys juice rewards instead (Figure 1A, right). After successfully clearing the maze, the monkeys would also receive additional rewards. The fewer attempts the animals made, the more rewards they would be given.

**Figure 1.**
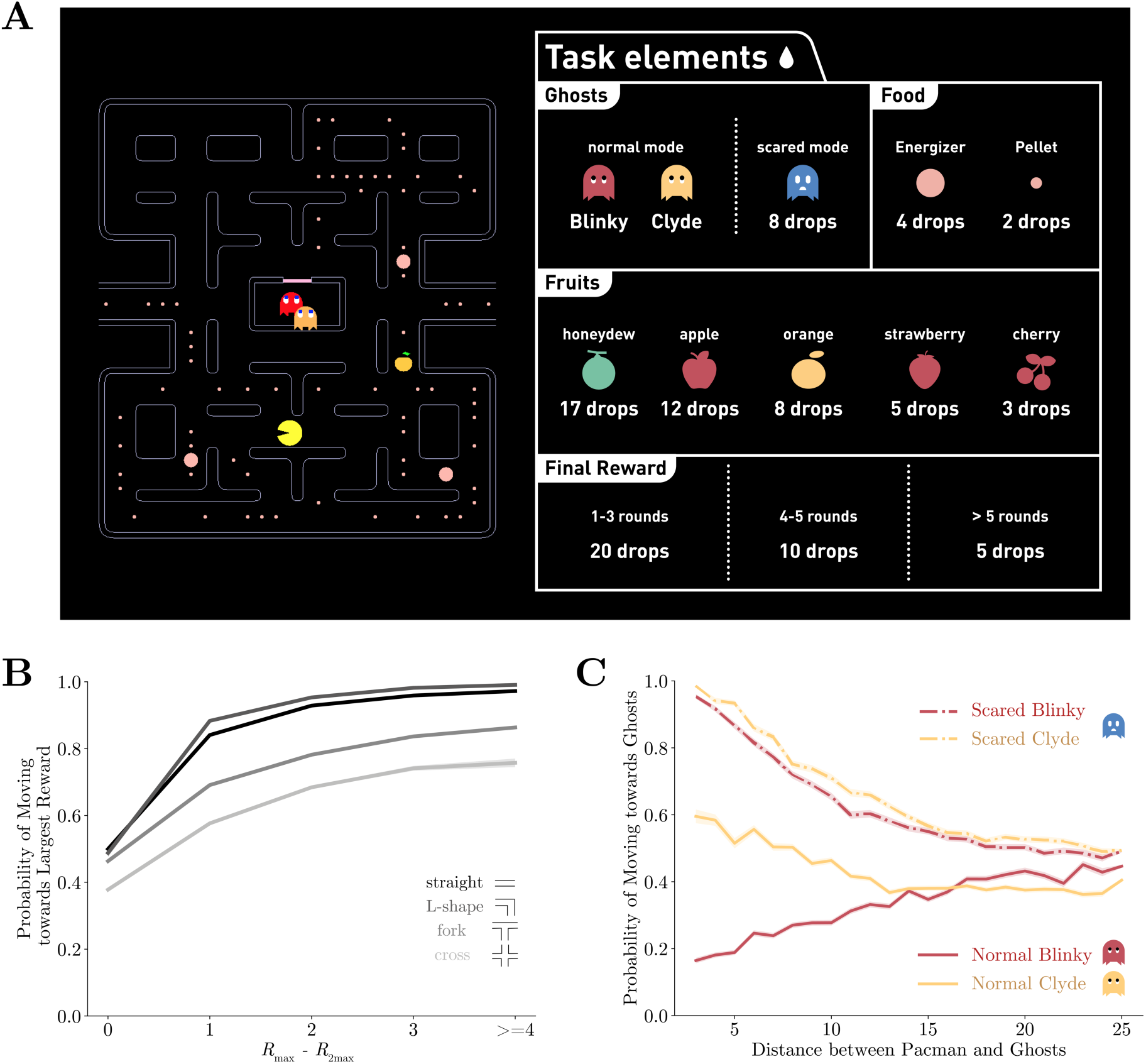
The Pac-Man game and the performance of the monkeys. **A**. The monkeys used a joystick to navigate Pac-Man in the maze and collect pellets for juice rewards. Also in the maze, there were two ghosts, Blinky and Clyde. The maze was fixed, but the pellets and the energizers were placed randomly initially in each game. Eating energizers turned the ghosts into the scared mode for 14 seconds, during which they were edible. There were also fruits randomly placed in the maze. The juice rewards corresponding to each game element are shown on the right. **B**. The monkeys were more likely to move toward the direction with more local rewards. The abscissa is the reward difference between the most and the second most rewarding direction. Different grayscale shades indicate path types with different numbers of moving directions. **C**. The monkeys escaped from the normal ghosts and chased the scared ghosts. The abscissa is the Dijkstra distance between Pac-Man and the ghosts. Dijkstra distance measures the distance of the shortest path between two positions on the map.

The game was essentially a foraging task for the monkeys. The maze required navigation, and to gain rewards, the animals had to collect pallets with the risk of encountering predators. Therefore, the game was intuitive for the monkeys, which was crucial for the training success. The training started with simple mazes with no ghosts, and more elaborated game elements were introduced one by one throughout the training process (see Methods and Figure S1 for detailed training procedures).

The behavior analyses include data from 74 testing sessions after all the game elements were introduced to the monkeys and their performance reached a reasonable level. We recorded the joystick movements, eye movements, and pupil sizes of the animals during the game. On average, the animals completed 33±9 (mean±SE) games in each session and each game took them 4.9±1.8 attempts (Figure S2).

Optimal gameplay requires the monkeys to consider a large number of factors, and many of them vary throughout the game either according to the game rules or as a result of the monkeys’ own choice. Finding the optimal strategy poses a computational challenge not only for monkeys but also for human and AI agents alike. The monkeys learned the task and played game well, as one can see from the example games (Movie S1-S3 for Monkey O, Movie S4-S5 for Monkey P). As a starting point to understand how monkeys solve the task, we first studied if the monkeys understand the basic game elements, namely, the pellets and the ghosts.

First, we analyzed the monkeys’ decision-making concerning the local rewards, which included the pellets, the energizers, and the fruits, within five tiles from Pac-Man for each direction. The monkeys tended to choose the direction with the largest local reward (Figure 1B). The probability of choosing the most rewarding direction decreased with the growing number of available directions, suggesting a negative effect of option numbers on the decision-making optimality.

The monkeys also understood how to react to the ghosts in different circumstances. The likelihood of the Pac-Man moving toward or away from the ghosts in different modes was plotted in Figure 1C. As expected, the monkeys tended to avoid the ghosts in the normal mode and chase them when they were scared. Interestingly, the monkeys picked up the subtle difference in the ghosts’ “personalities”. By design, Blinky aggressively chases Pac-Man, but Clyde avoids Pac-Man when they get close (See Methods). Accordingly, both monkeys were more likely to run away from Blinky but ignored Clyde even when it was close by. The ghosts were treated the same by the monkeys when in scared mode. The monkeys went after the scared ghosts when they were near Pac-Man.

These analyses suggest that the monkeys understood the basic elements of the game. While they reviewed some likely strategies of the monkeys, collecting local pellets and escaping or eating the ghosts, they only explained some of the monkeys’ decision making. Many other factors as well as the interaction between them affected the monkeys’ decisions. More sophisticated behavior was required for optimal performance, and to this end, the dynamic compositional strategy model was developed to understand the monkeys’ behavior.

### Basis strategies

While the overall goal of the game is to clear the maze, the monkeys may adopt different strategies for smaller objectives in different circumstances. We use the term “strategy” to refer to the solution for these sub-goals, and each strategy involves a smaller set of game variables with easier computation for decision making that forms actions.

We considered six intuitive and computationally simple strategies as the basis strategies. The *local* strategy moves Pac-Man toward the direction with the largest reward within 10 tiles. The *global* strategy moves Pac-Man toward the direction with the largest overall reward in the maze. The *energizer* strategy moves Pac-Man toward the nearest energizer. The two *evade* strategies move Pac-Man away from Blinky and Clyde in the normal mode, respectively. Finally, the *approach* strategy moves Pac-Man toward the nearest ghost. At any time during the game, monkeys could adopt one or a mixture of multiple strategies for decision making. These basis strategies, although not necessarily orthogonal to each other, can be linearly combined to explain the monkeys’ behavior.

At any time during the game, monkeys could adopt one or a mixture of multiple strategies for decision making. We assumed that the final decision for Pac-Man’s moving direction was based on a linear combination of the basis strategies, and the relative strategy weights were stable for a certain period. We used a change-point detection algorithm to segment each trial into such segments, then estimate the strategy weights by pooling the predictions from each strategy to match the monkeys’ actual choices.

### Monkeys adopt different strategies at different game stages

In Figure 2A, we plotted the normalized strategy weights in an example game segment (Movie S6). In this example, the monkey started with the *local* strategy and grazed pellets. With the ghosts getting close, it initiated the *energizer* strategy and went for a nearby energizer. Once eating the energizer, the monkey switched to the *approach* strategy to hunt the scared ghosts. Afterward, the monkey resumed the *local* strategy and then used the *global* strategy to navigate towards another patch when the local rewards were depleted. The dynamic compositional strategy model (See Methods for details) faithfully describes the monkey’s behavior by explaining Pac-Man’s movement with an accuracy of 0.971 in this example.

**Figure 2.**
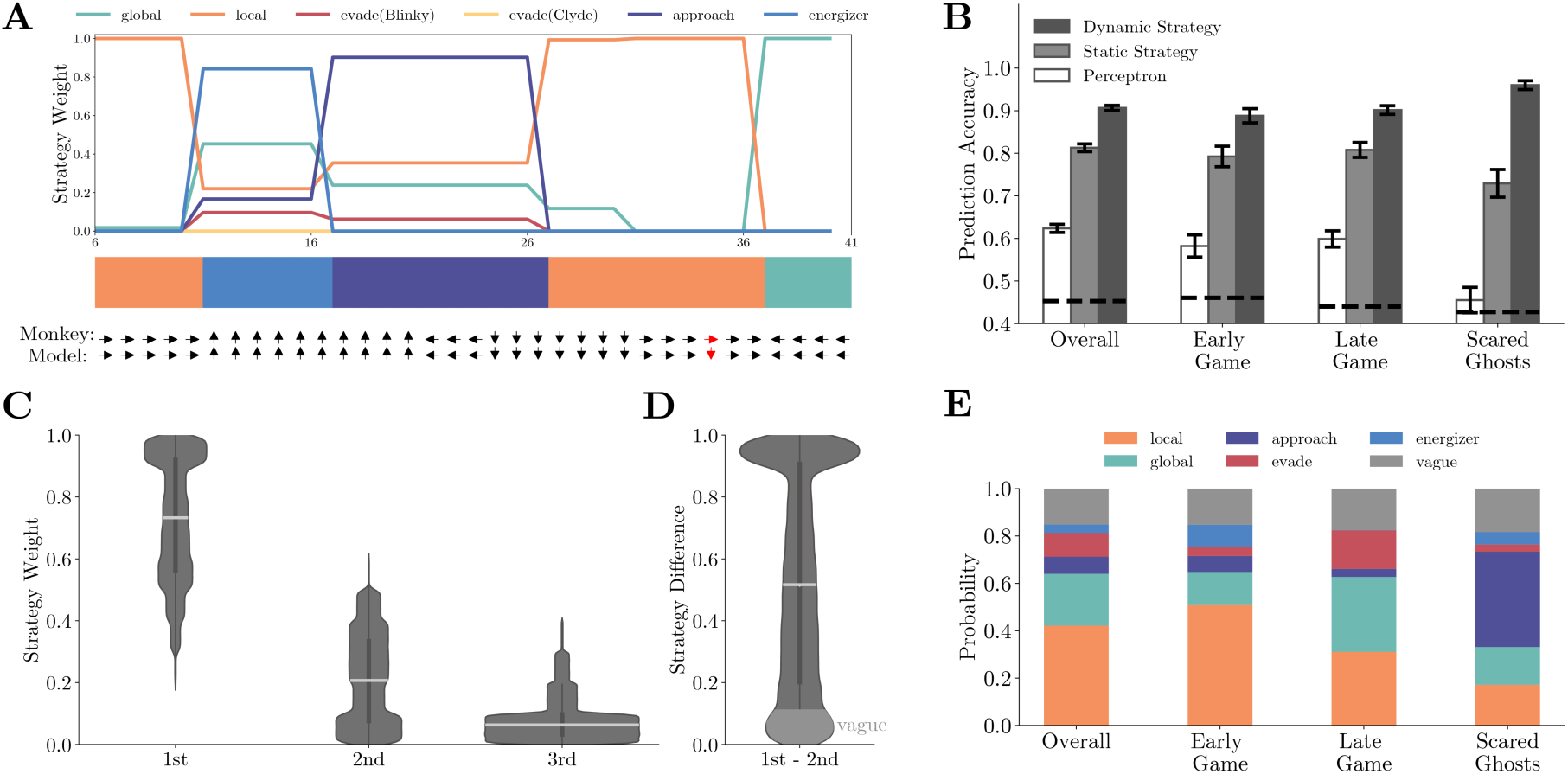
Fitting behavior with basis strategies. **A**. The normalized strategy weights in an example game segment. The horizontal axis is the time step. Each time step is 417 ms, which is the time that it takes Pac-Man to move across a tile. The color bar indicates the dominant strategies across the segment. The monkey’s actual choice and the corresponding model prediction at each time step are shown below, with red indicating a mismatch. The prediction accuracy for this segment is 0.971. Also, see Movie S6. **B**. Comparison of prediction accuracy across three models in four game contexts. See Table S3 and S4 for details. **C**. The histograms of the three dominating strategies’ weights. The most dominating strategy’s weights (0.736±0.209) were significantly larger than the secondary strategy (0.201±0.156) and tertiary strategy (0.051±0.079) by far. **D**. The histogram of the weight difference between the most and the second dominating strategies. The distribution is heavily skewed toward 1. In over 85% of the time, the weight difference was larger than 0.1, and more than a quarter of the time the difference was over 0.9. **E**. The ratios of labeled dominating strategies across four game contexts. In the early game, the *local* strategy was the dominating strategy. In comparison, in the late game, both the *local* and the *global* strategies had large weights. The weight of the *approach* strategy was largest when the ghosts were in the scared mode.

Overall, the dynamic compositional strategy model explains the monkeys’ behavior well. The model’s prediction accuracy is 0.910±0.015 for monkey O and 0.904±0.008 for monkey P. In comparison, a static strategy model, which uses the fixed strategy weights, achieves an overall accuracy of 0.804±0.015 and 0.822±0.011 for monkeys O and P, respectively (Figure 2B). The static strategy model’s accuracy still seems high, reflecting the fact that the monkeys were occupied with collecting pellets most of the time in the game. Thus, a combination of local and global strategies was often sufficient for explaining the monkeys’ choice. However, the average accuracy measurement alone and the fixed model could not reveal the monkeys’ adaptive behavior. The strategy dynamics are evident when we look at different game situations (Figure 2B). During the early game, defined as when there were more than 90% remaining pellets in the maze, the *local* strategy dominated all other strategies. In comparison, during the late game, defined as when there were fewer than 10% remaining pellets, both the *local* and the *global* strategies had large weights. The *approach* strategy came online when one or both scared ghosts were within 10 tiles around Pac-Man. The model’s prediction accuracies for the early game, late game, and scared-ghosts situation were 0.889±0.017, 0.902±0.010, and 0.960±0.010, which were significantly higher than the static strategy model’s accuracies (early: 0.792±0.024, p<0.001, late: 0.808±0.018, p<0.01, scared ghosts: 0.729±0.033, p<10^−6^; two-sample t-test).

The dynamic compositional strategy decision-making model is hierarchical. Each strategy produces a Pac-Man movement decision, and the decisions from the strategies are pooled to produce the final decision. It contrasts with flat decision-making models, in which Pac-Man movement decisions are computed directly from the game states. The latter is more computationally challenging. We trained a perceptron network as a representative flat model. The perceptron had one layer of 64 neurons for Monkey P and 16 neurons for Monkey O (The hyperparameters were determined by the highest fitting accuracy with 5-fold cross-validation, see Methods for details). The inputs were the same features used in our six strategies, and the outputs were the joystick movements. The perceptron model could only achieve 0.624±0.010 overall prediction accuracy (Figure 2B, white bars) and performed worse under each game situations (early: 0.582±0.026, p<10^−40^, late: 0.599±0.019, p<10^−16^, scared ghosts: 0.455±0.030, p<10^−16^; two-sample t-test).

### Monkeys adopted Take-the-best (TTB) heuristic

The model or the fitting procedure does not limit the number of strategies that may contribute to the monkeys’ choices at any time, yet the fitting results show that a single strategy often dominated the monkeys’ behavior. In the example (Figure 2A, Movie S6), the monkey switched between different strategies with one dominating strategy at each time point. This was a general pattern. We ranked the strategies according to their weights at each time point. The histograms of the three dominating strategies’ weights from all time points show that the most dominating strategy’s weights (0.736±0.209) were significantly larger than those of the secondary strategy (0.201±0.156) and tertiary strategy (0.051±0.079) by a significant extent (Figure 2C). The weight difference between the first and the second most dominating strategies was heavily skewed toward one (Figure 2D). These results are consistent with the Take-the-best heuristics in which using a single strategy and focusing on a particular aspect of the game at a time simplifies decision making.

Therefore, we labeled the monkeys’ strategy at each time point with the dominating strategy. When the weight difference between the dominating and secondary strategies was smaller than 0.1, the strategy was labeled as “*vague*”. The *local* and the *global* strategy were most frequently used overall. The *local* strategy was particularly prevalent during the early game when the local pellets were abundant, while the *global* strategy contributed significantly during the late game when the local pellets were scarce (Figure 2E).

### Strategy manifested in behavior

The strategy fitting procedure is indifferent to how the monkeys chose between the strategies, but the fitting results provide us some hints. The probability of the monkeys adopting the *local* or the *global* strategy correlated with the availability of local rewards: abundant local rewards lead to the *local* strategy (Figure 3A). On the other hand, when the ghosts were scared, the decision between chasing the ghosts and going on collecting the pellets depended on the distance between Pac-Man and the scared ghosts (Figure 3B). In addition, during the *global* strategy, the monkeys often moved Pac-Man to reach a patch of pellets far away from its current location. They chose the shortest path (Figure 3C) and made the fewest turns to do so (Figure 3D), demonstrating their goal-directed path-planning behavior under the particular strategy.

**Figure 3.**
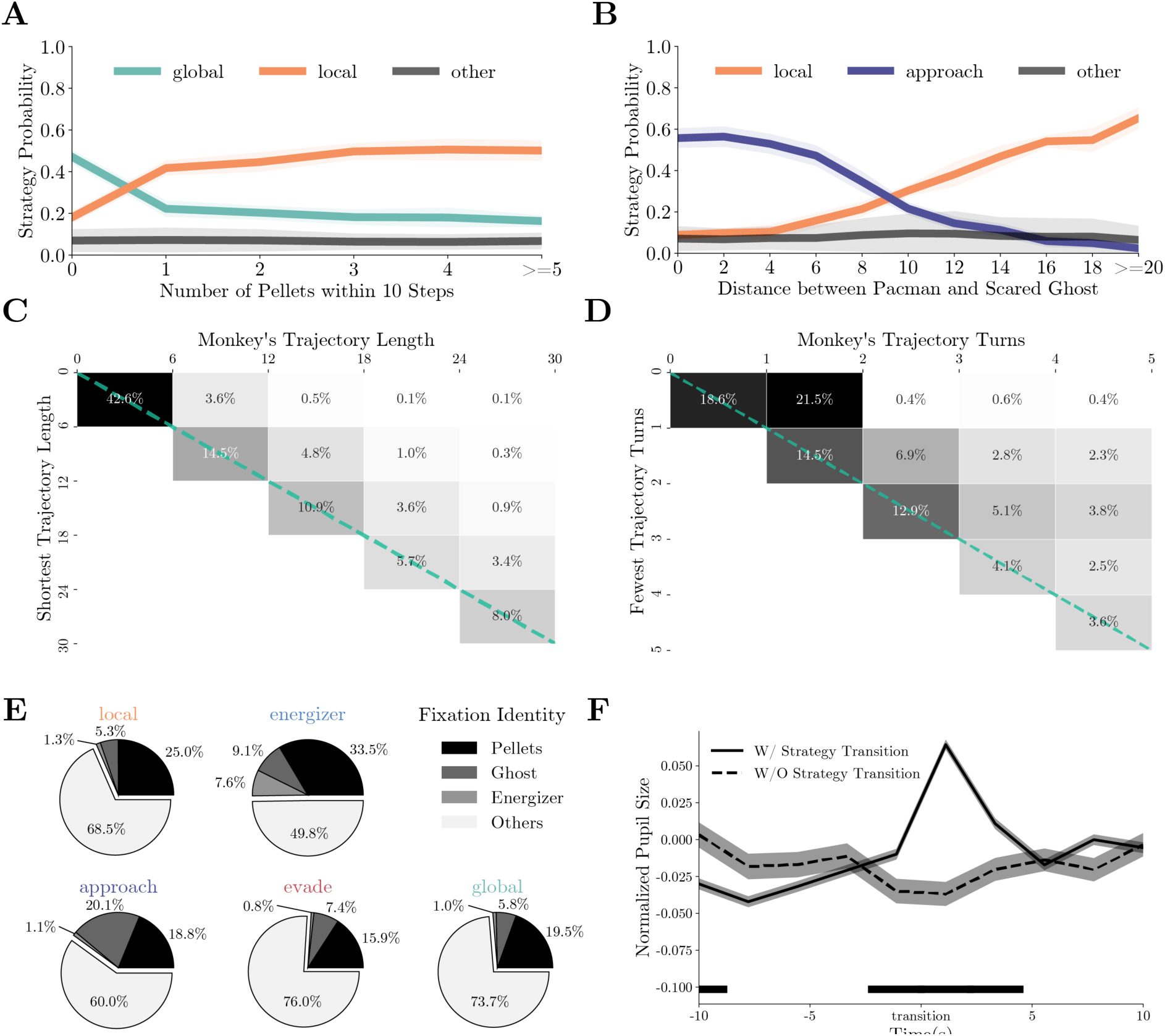
Monkeys’ behavior under different strategies. **A**. The probabilities of the monkeys adopting the *local* or *global* strategy correlate with the number of local pellets. **B**. The probabilities of the monkeys adopting the *local* or *approach* strategy correlate with the distance between Pac-Man and the ghosts. **C**. When adopting the *global* strategy to reach a far-away patch of pellets, the monkeys’ actual trajectory length was close to the shortest. The column denotes the actual length, and the row denotes the optimal number. The percentages of the cases with the corresponding actual lengths are presented in each cell. High percentages in the diagonal cells indicate close to optimal behavior. **D**. When adopting the *global* strategy to reach a far-away patch of pellets, the monkeys’ number of turns was close to the fewest possible turns. The column denotes the actual turns, and the row denotes the optimal number. The percentages of the cases with the corresponding optimal numbers are presented in each cell. High percentages in the diagonal cells indicate close to optimal behavior. **E**. Average fixation ratios of ghosts, energizers, and pellets when the monkeys used different strategies. **F**. The monkeys’ pupil diameter increase around the strategy transition (solid line). Such increase was absent if the strategy transition went through the *vague* strategy (dashed line). Black bar at the bottom denotes p < 0.01, two-sample t-test.

The fitting results can be further corroborated from the monkeys’ eye movements and pupil dilation. Because different game aspects were used in different strategies, the monkeys should be looking at different things when using different strategies. We classified monkeys’ fixation locations into three categories: ghosts, energizers, and pellets (See Methods for details). Figure 3E shows the fixation ratio of these game objects under different strategies. Although a large number of fixations were directed at the pellets in all situations, they were particularly frequent under the *local* and *energizer* strategies. Fixations directed to the energizers were scarce, unless when the monkeys adopted the *energizer* strategy. On the other hand, monkeys looked at the ghosts most often when the monkeys were employing the *approach* strategy to chase the ghosts (p < 0.001, two-sample t-test). Interestingly, the monkeys also looked at the ghosts more often under the *energizer* strategy than under the *local* strategy (p < 0.001, two-sample t-test), which suggests that the monkeys were also keeping track of the ghosts when going for the energizer.

While the fixation patterns revealed that the monkeys paid attention to different game elements in different strategies, we also identified a physiological marker that reflected the strategy switches in general but was not associated with any particular strategy. Previous studies revealed that non-luminance-mediated changes in pupil diameter can be used as markers of arousal, surprise, value, and other factors during decision making (Joshi & Gold, 2020). Here, we analyzed the monkeys’ pupil dilation during strategy transitions. When averaged across all types of transitions, the pupil diameter exhibited a significant but transient increase around strategy transitions (p < 0.01, two-sample t-test, Figure 3F). Such an increase was absent when the strategy transition went through a *vague* period. This increase was evident in the transitions in both directions, e.g., from *local* to *global* (Figure S6A, S6C) and from *global* to *local* (Figure S6B, S6D). Therefore, it cannot be explained by any particular changes in the game state, such as the number of local pellets. Instead, it reflected an internal state change during strategy switches.

### Compound strategies

The compositional strategy model divides the monkeys’ decision making into different hierarchies (Figure 4). At the lowest level, decisions are made for the motor movements: up, down, left, or right. The basis strategies in our model provide a heuristic for the monkey to reduce the complexity of this decision making by only considering a subset of game parameters. At the middle level, choices are made between these basis strategies. The pupil dilation change reflected the decision making at this level. At a higher level, the basis strategies may be pieced together for more sophisticated compound strategies. These compound strategies are not simple impromptu strategy assemblies. Instead, they may reflect more advanced planning. During the gameplay, monkeys may initiate a compound strategy to deal with a particular situation. Here, building on the strategy analyses, we describe two scenarios in which we can see this behavior.

**Figure 4.**
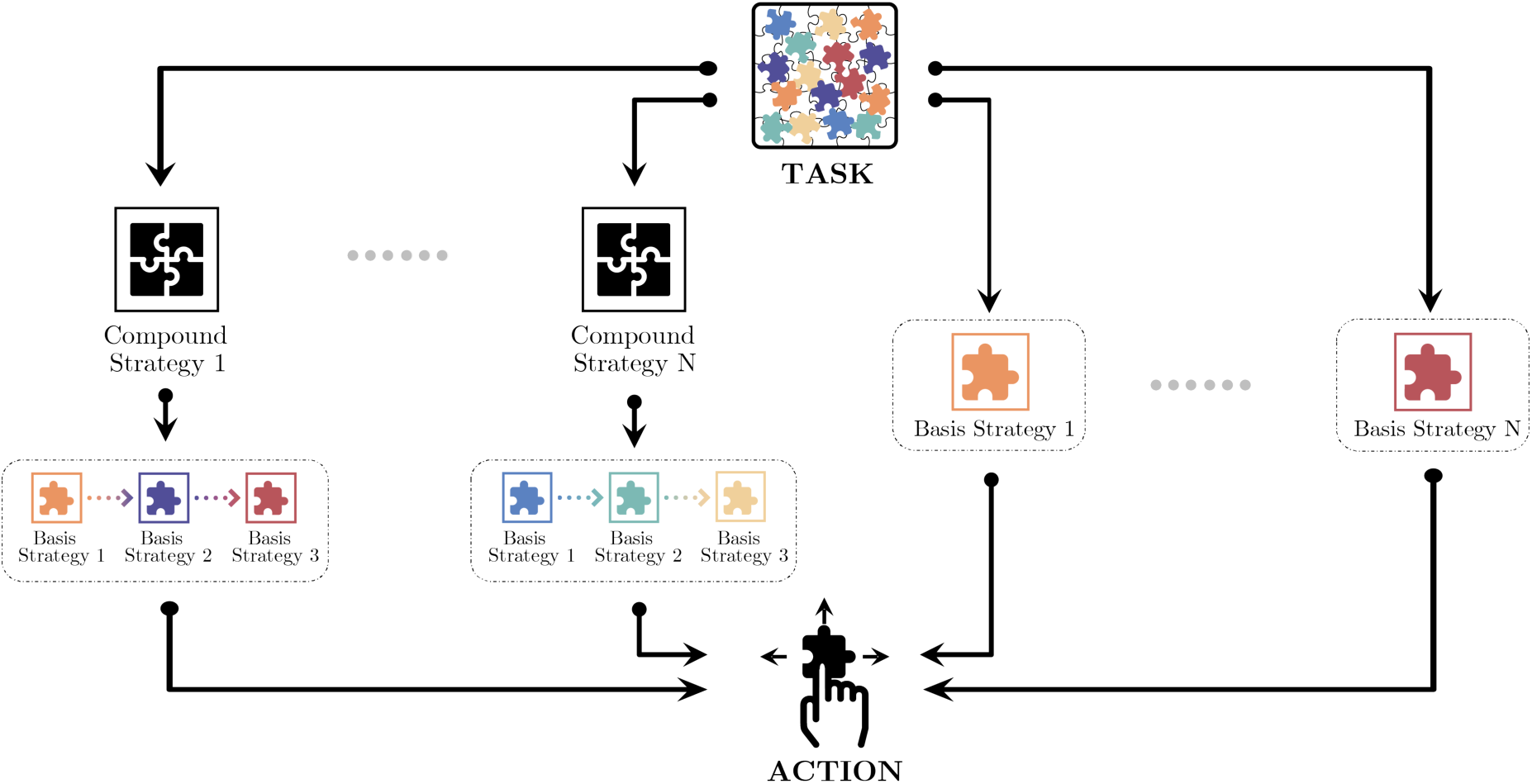
Monkeys’ decision making in different hierarchies. At the lowest level, decisions are made for the motor movements: up, down, left, or right. At the middle level, choices are made between the basis strategies. At a higher level, the simple strategies may be pieced together for more sophisticated compound strategies.

The first scenario involves the energizers, which is an interesting feature of the game. They not only provide an immediate reward but also lead to potential future rewards from eating ghosts. With the knowledge that the effect of the energizers was only transient, the monkeys could plan accordingly to maximize their gain from the energizers. In some trials, the monkeys immediately switched to the *approach* strategy after eating an energizer (Movie S7). In contrast, sometimes the monkey appeared to treat an energizer just as a more rewarding pellet. They continued collecting pellets with the *local* strategy after eating the energizer and might happen to catch a ghost on the way (Movie S8). Accordingly, we distinguished these two behaviors using the strategy labels after the energizer consumption and named the former as *planned attack* and the latter as *accidental consumption* (See Methods for details). With this criterion, we extracted 493 (Monkey O) and 463 (Monkey P) *planned attack* plays, and 1970 (Monkey O) and 1295 (Monkey P) *accidental consumption* plays in our data set.

The strategy weight dynamics around the energizer consumption showed distinct patterns when the animals adopted the compound strategy *planned attack* (Figure 5A). When the monkeys carried out *planned attacks*, they started to approach the ghosts well before the energizer consumption, which is revealed by the larger weights of the *approach* strategy than that in the *accidental consumption*. The monkeys also cared less for the local pellets in *planned attacks* before the energizer consumption. The weight dynamics suggest that the decision of switching to the *approach* strategy was not an afterthought but planned well ahead. The monkeys strung the *energizer*/*local* strategy with the *approach* strategy together into the compound strategy to eat an energizer and then hunt the ghosts. Such a compound strategy should only be employed when Pac-Man, ghosts, and an energizer are in close range. Indeed, the average of the distance between the Pac-Man, the energizer, and the ghosts was significantly smaller in *planned attack* than in *accidental consumption* (Figure 5B, p < 0.001, two-sample t-test).

**Figure 5.**
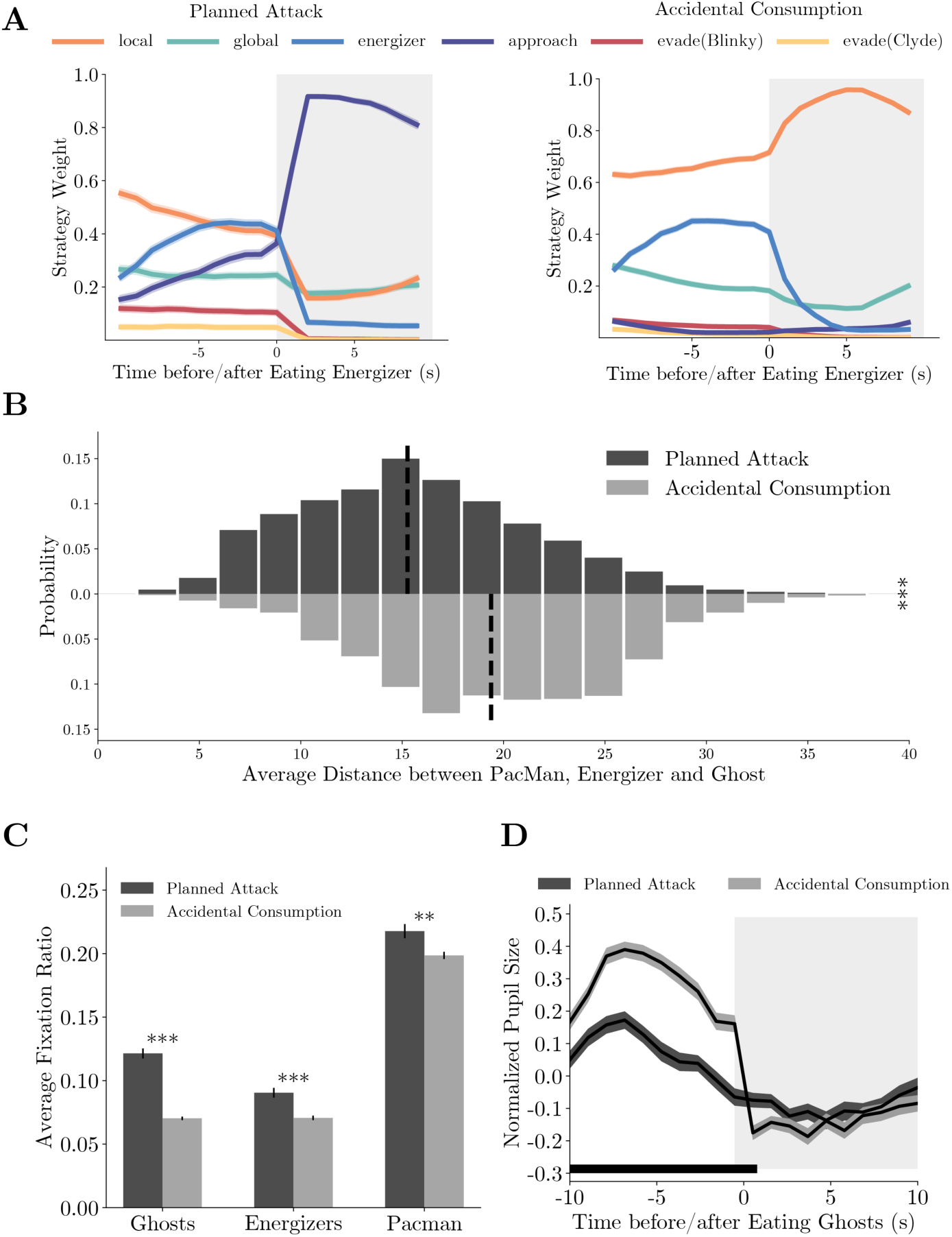
Compound strategies: *planned attack*. **A**. Average strategy weight dynamics in *planned attacks* (left) and *accidental consumptions* (right). **B**. The average distance between the Pac-Man, energizer, and the ghosts in *planned attacks* and *accidental consumptions*. *** denotes p<0.001, two-sample t-test. **C**. Ratios of fixations on the ghosts, the energizer, and Pac-Man. *** denotes p<0.001 and ** denotes p<0.01, two-sample t-test. **D**. The pupil size aligned to the ghost consumption. The black bar near the abscissa denotes data points where the two traces are significantly different (p < 0.01, two-sample t-test).

Again, the *planned attacks* were also associated with distinct eye movement and pupil size dynamics. The monkeys fixated on the ghosts, the energizers, and Pac-Man more frequently before the energizer consumption in *planned attacks* than in *accidental consumption* (Figure 5C, p<0.001, two-sample t-test), reflecting more active planning under the way. In addition, the monkeys’ pupil sizes were smaller before they caught a ghost in *planned attacks* than in *accidental consumption* (p < 0.01, two-sample t-test), which may reflect a lack of surprise under *planned attacks* (Figure 5D). The difference was absent after the ghost was caught.

The second scenario involves a counter-intuitive move in which the monkeys move Pac-Man towards a normal ghost to be eaten on purpose. Although the move appeared to be suboptimal, it was beneficial in a certain context. The death of Pac-Man resets the game and returns Pac-Man and the ghosts to their starting positions in the maze. As the only punishment in the monkey version of the game is a timeout, it is advantageous to reset the game by committing such suicide when local pellets are scarce, and the remaining pellets are far away.

To analyze this behavior, we defined the compound strategy *suicide* using strategy labels. We computed the distances between Pac-Man and the closest pellets before and after its death. In *suicides*, the Pac-Man’s death significantly reduced this distance (Figure 6A upper histogram, Movie S9). This was not true when the monkeys were adopting the *evade* strategy but failed to escape from the ghosts (*failed evasions*, Figure 6A bottom histogram, Movie S10). In addition, the distance between Pac-Man and the ghosts was greater in *suicides* (Figure 6B, p < 0.001, two-sample t-test). Therefore, these suicides were a proactive decision. Consistent with the idea, the monkeys tended to saccade toward the ghosts and pellets in *suicides* (Figure 6C, p < 0.001, two-sample t-test). Their pupil size decreased even before the Pac-Man’s death in *suicides*, which was significantly smaller than in *failed evasions*, suggesting that the death was anticipated (Figure 6D, p < 0.01, two-sample t-test).

**Figure 6.**
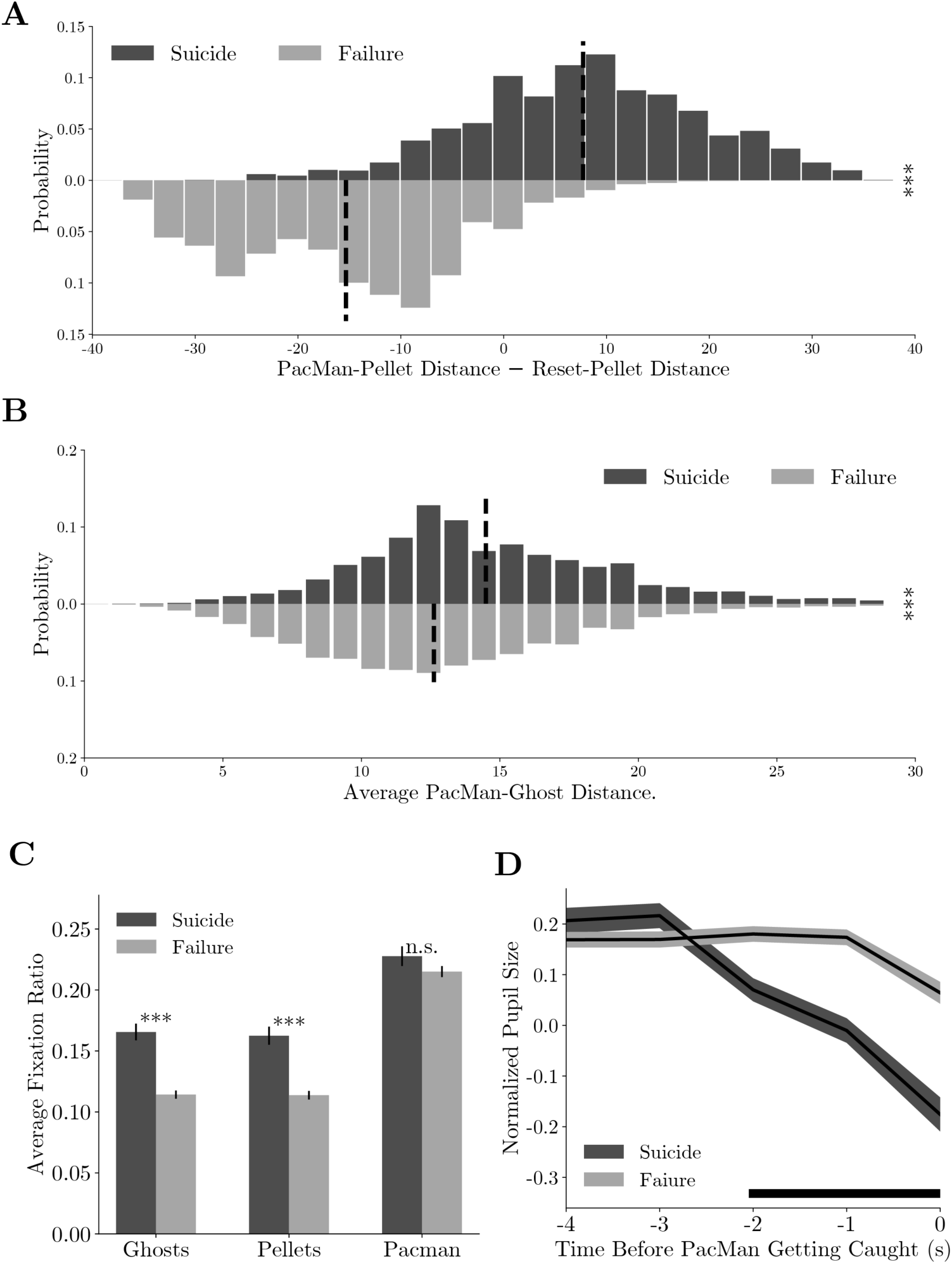
Compound strategies: *suicide*. **A**. Distance difference between Pac-Man and closest pellet before and after the death are smaller in *suicides* than in *failed evasions*. *** denotes p<0.001, two-sample t-test. **B**. Average distance between Pac-Man and the ghosts was greater in *suicides* than in *failed evasions*. *** denotes p<0.001, two-sample t-test. **C**. The monkeys fixated more frequently on the ghosts and the pellets in *suicides* than in *failed evasions*. *** denotes p<0.001, two-sample t-test. **D**. The monkeys’ pupil size decreased before the Pac-Man’s death in *suicides*. The black bar near the abscissa denotes data points where the two traces are significantly different (p < 0.01, two-sample t-test).

Together, these two examples demonstrated that how monkeys’ advanced game play can be understood with concatenated basis strategies. The compositional strategy model not only provides a good fit for the monkeys’ behavior but also offers insights into the monkeys’ gameplay.

## Discussion

Just as one cannot gain a full understanding of the visual system by studying it using bars and dots, pursuing a deeper insight into the cognitive capability of the brain demands sophisticated behavior paradigms in which an ensemble of perception, attention, valuation, executive control, decision-making, motor planning, and other cognitive processes need to work together continuously across time. Naturally, quantifying and modeling these behaviors in such paradigms is challenging, but here we demonstrated that the behavior of monkeys during a complex game can be understood and described with a set of basis strategies that decompose the decision making into different hierarchies.

Although the particular set of basis strategies in the model were hand-crafted, we have good reasons to believe that they reflect the decision-making process from the monkey. First, the model fitting procedure is agnostic to how one should choose between the strategies, yet the resulting strategies can be corroborated both from monkeys’ route planning, eye movements, and pupil dilation patterns. This is evidence for both the validity of the model and the rationality behind monkeys’ behavior. The correlation between the results of strategy fitting and the fixation patterns of the monkeys indicates that the animals learned to selectively attend to the features that were relevant for their current strategy while ignoring others to reduce the cognitive load for different states. Similar behaviors have also been observed in human studies (Leong et al., 2017; Wilson & Niv, 2012). In addition, the pupil dilation pattern, which cannot be explained by any sensory or reward events related to any particular strategies, was an indication of the extra cognitive processing carried out in the brain to handle the strategy transitions. Finally, the model, without being specified so, revealed that a single strategy dominates monkeys’ decision-making during most of the game. This is consistent with the idea that the strategy-using as the method that the brain uses to simplify decision making by ignoring irrelevant game aspects to solve complex tasks (Binz et al., 2020; Moreno-Bote et al., 2020).

In some previous animal studies, strategies were equated to decision rules (Bunge & Wallis, 2007; Genovesio & Wise, 2007; Hoshi et al., 2000; Mante et al., 2013; Tsujimoto et al., 2011). The rules were typically mutually exclusive, and the appropriate rule was either specified with explicit sensory cues or the trial-block structure. The rules in these studies can be boiled down to simple associations, even in cases when the association may be abstract (Genovesio & Wise, 2007). In the current study, however, we defined a set of strategies as a heuristic that reduces a complex computation into a set of smaller and more manageable problems or computations. There are no explicit cues or trial block structures to instruct animals on which strategies to choose. Nevertheless, the existing literature indicates that a broad prefrontal network, including the dorsolateral prefrontal cortex, orbitofrontal cortex, and polar cortex, is engaged in rule use and rule switching. This network is likely to play important roles in strategy-based decision making, too.

Our Pac-Man paradigm elicits monkeys’ more complex and natural cognitive ability. First, the game contains an extensive state space. This requires monkeys to simplify the task by developing temporally extended strategies to accomplish sub-goals. Second, there exists non-exclusive solutions or strategies to solve the Pac-Man task appropriately. Instead of spoon-feeding monkeys the exact solution in simple tasks, we trained them with all relevant game elements during the training phases and allowed them to proactively coordinate and select strategies freely. Therefore, our Pac-Man paradigm does not restrict monkeys’ behavior with a small number of particular rules and allows the brain and its neural circuitry to be studied in a more natural setting (Krakauer et al., 2017).

In summary, our model distilled a complex task into different layers of decision making centered around a set of compositional strategies, which paved the way for future experiments that will provide key insights into the neural mechanisms underlying sophisticated cognitive behavior that go beyond what most of the field currently studies.

## Acknowledgments

We thank Wei Kong, Lu Yu, Ruixin Su, Yunxian Bai, Zhewei Zhang, Yang Xie, Yiwen Xu, and Yue Hao for their help in all phases of the study, and Liping Wang and Xaq Pitkow for providing comments and advice.

## Author contributions

Conceptualization: TY

Methodology: QY, ZL, WZ, TY

Investigation: QY, ZL, WZ, JL, XC, JZ, TY

Visualization: QY, WZ, JL, JZ, TY

Funding acquisition: TY

Project administration: TY

Supervision: TY

Writing – original draft: QY, TY

Writing – review & editing: QY, ZL, WZ, JZ, TY

## Data and materials availability

The data that support the findings of this study are available from the corresponding author upon reasonable request.

## Supplementary Materials

Materials and Methods

Figs. S1 to S8

Tables S1 to S6

Captions for Movies S1 to S10

## Materials and Methods

### Subjects and Materials

Two male rhesus monkeys (Macaca mulatta) were used in the study (O and P). They weighed on average 6-7 kg during the experiments. All procedures followed the protocol approved by the Animal Care Committee of Shanghai Institutes for Biological Sciences, Chinese Academy of Sciences (Shanghai, China).

### Training Procedure

To help monkeys to understand Pac-Man game elements and develop their decision-making strategies, we divided the training procedures into the following three stages. In each stage, we gradually increased game depth based on their conceptual and implementational complexity.

#### Stage One: Reward

In the first stage, the monkeys were trained to use the joystick to control Pac-Man to navigate through simple mazes for consuming pellets (Figure S1A). Training began with horizontal and vertical linear mazes. In each maze, Pac-Man started from the center where pellets were at one end and a static ghost was at the other end. Monkeys earned 2 drops of juice (1 drop= 0.5 mL) immediately when consuming a pellet. Monkeys could earn an extra-large amount of juice by clearing all pellets. Running towards the static ghost would lead to the end of the trial with a time-out penalty (5 s). When the monkeys completed more than 100 correct trials with above 80% accuracy, we introduced two slightly more complex mazes, the T and the upside-down T maze. After the monkeys completed more than 50 correct trials in T-mazes with above 80% accuracy, we introduced the H-maze. Stage One training included 58 sessions for Monkey O and 84 sessions for Monkey P.

#### Stage Two: Ghost

In the second stage, the monkeys were trained to deal with the ghosts (Figure S1B). In addition, the mazes used in this stage were closed and had loops. A ghost would block one of the routes leading to the pellets, forcing the monkeys to take alternative routes. In the first phase, the ghost was stationary in a square maze. Pac-Man started from one of the four corners, and pellets were distributed in two adjacent arms. The ghost was placed at the corner where the two arms joined, forcing Pac-Man to retreat after clearing the pellets in one arm. In the second phase, the ghost moved within the arm. In the third phase, the ghost would chase Pac-Man. Stage Two training included 86 sessions for Monkey O and 74 sessions for Monkey P.

#### Stage Three: Energizer

In this stage, the monkeys were trained to understand the energizer (Figure S1C). In the first phase, the monkeys were trained to understand the distinction between normal and scared ghosts. We used the square maze with a normal ghost or a scared ghost randomly placed across trials. Blinky in the normal mode would chase Pac-Man, while in scared mode moved in a random direction at half of Pac-Man’s speed. Monkey earned 8 drops of juice after eating a scared ghost. In the second phase, the monkeys were trained with the maze that the scared mode could only be triggered by an energizer. Two energizers were randomly placed in each maze. Monkey earned 4 drops of juice when eating an energizer and turned ghosts into the scared mode immediately. The scared mode lasted 14 seconds. As a reminder, ghosts in the scared mode flashed for 2 seconds before turning back into normal mode. In the third phase, we adopted the maze used in our final game-play recording (Figure 1A). The detailed game rules can be found in the following Task Paradigm session. Stage Three training included 248 sessions for Monkey O and 254 sessions for Monkey P.

### Task Paradigm

The Pac-Man game in the current study is adapted from the original game by Namco®. All key concepts of the game are included. In the game, the monkey navigates a character named Pac-Man through a maze with a four-way joystick to collect pellets and energizers. The maze is sized at 700×900 pixels, displayed with the resolution of 1920×1080 on a 27-inch monitor placed at 68 cm away from the monkey. The maze can be divided into square tiles of 25×25 pixel^2^. The pellets and energizers are placed at the center of a tile and they are consumed when Pac-Man moves into the tile. In the recording sessions, there are 88 or 73 pellets in the maze, each worth 2 drops of juice, and 3 or 4 energizers, each worth 4 drops of juice. We divided the maze into 4 quarters, and the pellets and energizers are randomly placed in 3 of them, with one randomly chosen quarter empty. In addition, just as in the original game, there are five different fruits: cherry, strawberry, orange, apple, and melon. They are worth 3, 5, 8, 12, and 17 drops of juice, respectively. In each game, one of the fruits is placed at a random location at each game. The maze also contains two tunnels that teleport Pac-Man to the opposite side of the maze.

There are two ghosts in the game, Blinky and Clyde. They are released from the Ghost Home, which is the center box of the maze, at the beginning of each game. Blinky is red and is more aggressive. It chases Pac-Man all the time. Clyde is orange. It moves toward Pac-Man when it is more than 8 tiles away from Pac-Man. Otherwise, it moves towards the lower-left corner of the maze. The eyes of the ghosts indicate the direction they are traveling. The ghosts cannot abruptly reverse direction in the normal mode. Scared ghosts move slowly to the ghost pen located at the center of the maze. The scared state lasts 14 seconds and the ghosts flash as a warning during the last 2 seconds of the scared mode. Monkeys get 8 drops of juice if they eat a ghost. Dead ghosts move back to the ghost pen and then respawn. The ghosts can also move through the tunnels, but their speed is reduced when in the tunnel. For more explanations on the ghost behavior, please refer to https://gameinternals.com/understanding-pac-man-ghost-behavior.

When Pac-Man is caught by a ghost, it and the ghosts return to the starting location. When all the pellets and energizers are collected, the monkey receives a reward based on the number of rounds that takes the monkey to complete the game: 20 drops if the round number is from 1 to 3; 10 drops if the round number is from 4 to 5; 5 drops if the round number is larger than 5.

### Behavioral Data Recording and Preprocessing

We monitored the monkeys’ joystick movements, eye positions, and pupil sizes during the game. The joystick movements were sampled at 60Hz. We used Eyelink 1000 Plus to record two monkeys’ eye positions and pupil sizes. The sampling rate was 500Hz or 1000Hz.

The data we presented here are based on the sessions after the monkeys went through all the training stages and were able to play the game consistently. On average, monkeys completed 33±9 games in each session and each game took them 4.86±1.75 attempts. In total, we recorded 3217 games, 15772 rounds, and 899381 joystick movements. The monkeys’ detailed game statistics were shown in Figure S2.

### Basic Performance Analysis

In Figure 2B, we compute the rewards for each available moving direction at each location by summing up the pellets rewards (1 unit) and energizers rewards (2 units) within 5 steps from Pac-Man’s location. Locations are categorized into four path types defined in Table S1. For each path type, we calculate the probability that the monkey moved in the direction with the largest rewards conditioned on the reward difference between the most (*R*_*max*_) and the second most rewarding directions (*R*_2*max*_).

### Basis Strategies

We include six basis strategies in the dynamic strategy model. In each basis strategy, we compute the utility values for all directions (𝒟={left, right, up, down}), expressed as a vector with a length of four. Notice that not all directions are available, utility values for unavailable directions are set to zero. The moving direction is computed according to the largest average utility value for each strategy.

We determine the utility associated with each direction and its possible trajectories. Specifically, let *p* represents the Pac-Man’s position and *τ*(*p*) represent a path starting from p with the length of 10. We define *g* = {*gB, gC*} to be the position of two ghosts, Blinky and Clyde, and *r* = {*r*_*p*_, *r*_*e*_, *r*_*f*_} to be the positions of pellets, energizers, and fruits, respectively. We compute the utility of each path *τ*(*p*) as following (without specific noting, *τ*(*p*) is denoted as *τ* for simplicity).

We use the *local* strategy to describe the local graze behavior within a short distance with the utility function defined as

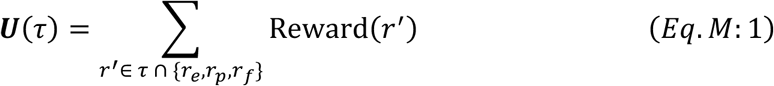

where *τ* ∩ {*r*_*e*_, *r*_*p*_, *r*_*f*_} denotes the pellets/energizers/fruits on the path. Specific parameters for awarded and penalized utilities of each game element in the model can be found in Table S2.

*Evade* strategy focuses on dodging close-by ghosts. Specifically, we create two evade strategies (*evade* Blinky and *evade* Clyde) that react to the respective ghost, with the utility function defined as

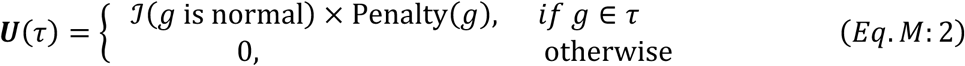

with *g* = *g*_*B*_ and *g* = *g*_*C*_ respectively. Here 𝒥(*s*) is an indication function where 𝒥(*s*) = 1 when statement *s* is true, otherwise 𝒥(*s*) = 0.

*Energizer* strategy moves Pac-Man toward the closest energizer. In this case, the rewards set *r* only contains the positions of energizers (i.e., *r* = {*r*_*e*_}):

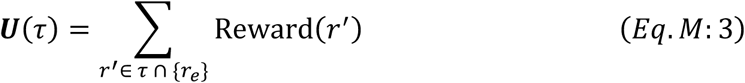

*Approach* strategy moves Pac-Man toward the ghosts, regardless of the ghosts’ mode. Its utility function is

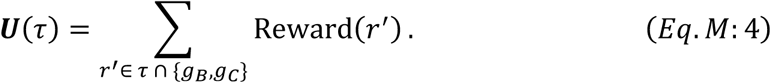

*Global* strategy does not use a decision tree. It counts the total number of pellets in the whole maze in each direction without considering any trajectories. For example, the utility for the down direction is the total number of pellets that sit vertically below the Pac-Man’s location.

We construct the utility of each agent as a vector *U*_*a*_ ∈ ℝ^4^, *f or a* ∈ {*local, global, evade Blinky, evade Clyde, approach, energizer*} of four directions. For each direction *d* ∈ 𝒟, its utility is obtained by averaging all the path utilities *U*(τ) in that direction, such as

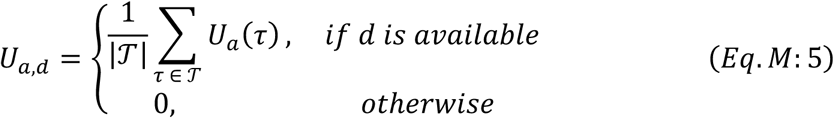

### Models and Model Fitting

We adopted a softmax policy and used a maximum likelihood estimate (MLE) to fit the model to the behavior.

#### Utility Preprocessing

To combine the strategies and produce a decision, we first preprocess the utility data computed from decision trees with two steps. First, because two *evade* strategies have negative utility values, we calculate their difference to the worst-case scenario within a trial and use the difference, which is a positive value, as the utility for the two *evade* strategies:

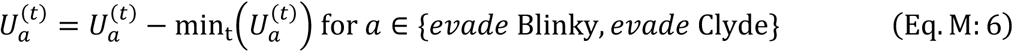

Second, because the scale of utility value varies in different strategies, we normalize the utilities within each strategy:

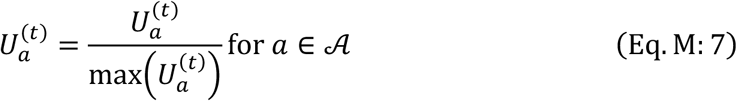

with 𝒜 = {*local, global, evade Blinky, evade Clyde, approach, energizer*}.

#### Softmax Policy

With the adjusted and normalized utility values, each strategy *a* is associated with a set of utility values ***U***_*a,d*_ for four directions *d* ∈ 𝒟. We compute the utility for each direction *d* by simply combining them linearly with strategy weights ***w*** ∈ ℝ^6^:

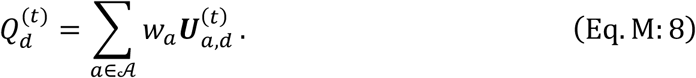

The final decision is based on a softmax policy:

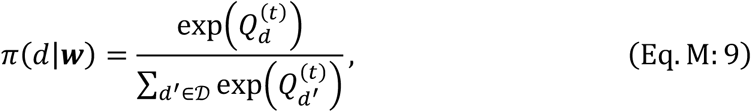

where *π*(*d*|***w***) describes the probability of choosing *d* given weights ***w***.

#### Maximum Likelihood Estimate

We used the Maximum Likelihood Estimate (MLE) approach to estimate monkeys’ strategy weights **w** in a time window *δ*. Based on Pac-Man’s actual moving directions ***d***^∗^, we computed the likelihood as

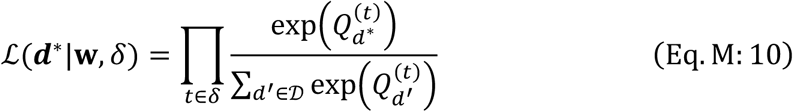

The strategy weights within a time window can be estimated by maximizing the log-likelihood:

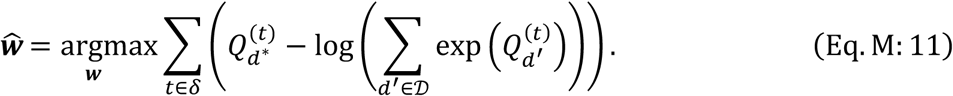

#### Dynamic Compositional Strategy Model

The dynamic compositional strategy model estimates the strategy weights using time windows of flexible length. We assumed that the relative strategy weights were stable for a period. The weights can be estimated from the monkeys’ choices during this period. We designed a two-pass fitting procedure to divide each trial into segments of stable strategies and extract the strategy weights for each segment, avoiding potential overfitting caused by segmentations too fine with too many weight parameters while still capturing the strategy dynamics. The procedure is as follows.

1. We first formulated fine-grained time windows Δ = {*δ*_1_, *δ*_2_, ⋯, *δ*_*K*_} according to the following events: Pac-Man direction changes, ghost consumptions, and energizer consumptions. The assumption is that the strategy changes only occurred at those events.
2. The first-pass fitting was done using the fine-grained time windows to get the maximum likelihood estimates of the strategy weights as a time series of 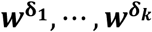
3. We then used a change-point detection algorithm to detect any changes in the strategy weights in the time series of 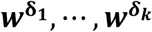. Specifically, we selected a changing points number *K* and used a forward dynamic programming algorithm (Truong et al., 2020) to divide the series into *K* segments *Δ*_*K*_ = {*δ*_1_, *δ*_2_, ⋯, *δ*_*K*_} by minimizing the quadratic loss 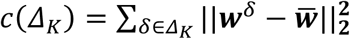. Here, 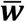 is the empirical mean of these fine-grained weights corresponding to segment sets, *Δ*_*K*_. With *Δ*_*K*_, we constructed the coarse-grained time windows by combining the fine-grained time windows.
4. The second-pass fitting was then done using the coarse-grained time windows *Δ*_*K*_ with MLE, and the sum of log-likelihood 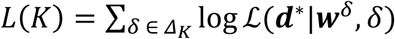 was the loss function.
5. We repeated the steps 3 and 4 with hyperparameter *K* tranversing through {2,3, ⋯, 20} to find out *K*^∗^ = *argmax L*(*K*). The final fitting results were based on the normalized fitted weights with *K*^∗^ coarse-grained time windows 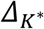.

#### Static Strategy Model

The static strategy model uses all data to estimate a single set of strategy weights.

#### Linear Perceptron Model

We built a linear perceptron as a representative flat descriptive model (without calculating utilities and strategies) to describe monkeys’ decision-making based on the same 20 task features used in the strategy model. These features include the modes of Blinky and Clyde (2 features), Dijkstra distances between Blinky and Pac-Man in four directions (4 features), Dijkstra distances between Clyde and Pac-Man in four directions (4 features), Dijkstra distances between the closest energizer and Pac-Man in four directions (4 features), distances between fruits and Pac-Man in four directions (4 features), the number of pellets within 10 steps of Pac-Man, and the number of pellets left in the maze. For unavailable directions, the corresponding feature value is filled with a None value. We trained a three-layer perceptron with monkeys’ choice behavior: an input layer for 20 features, a hidden layer, and an output layer for four directions. We used scikit-learn (https://scikit-learn.org/). The size of the hidden layer was selected from *n*_*hidden*_ ={16, 32, 64, 128, 256} with the largest average prediction accuracy on all data. For Monkey O, the best hidden unit number is 64, and for Monkey P, the best hidden unit number is 128. Each model used Adam for optimization, training batch size = 128, learning rate = 0.001, regularization parameter = 0.0001, and the activation function *f*(⋅) for the hidden layer was an identity function.

#### Model Comparison

We compared three models (static strategy model, dynamic strategy model, and linear perceptron model) in four game contexts shown in Table S3. We used 5-fold cross-validation to evaluate the fitting performance of these three models with each monkeys’ behavior data.

#### Strategy Heuristic Analysis

We label the behavior strategy as “*vague*” when the weight difference between the largest and the second largest strategies is less than 0.1 (Figure 2D). Otherwise, the label was based on the strategy with the largest weight.

In Figure 3A, B, we evaluated the strategy probability dynamics with respect to two features: Local pellet density (the number of pellets within 10 steps from Pac-Man) and Scared ghost distance. We grouped the behavior data based on these two features and calculated the frequency of the relevant strategies in each. Means and standard deviations were computed by bootstrapping 10 times with a sample size of 100 for each data point (Figure 3A, B).

In Figure 3C, D, Monkeys’ moving trajectories with at least four consecutive steps labeled as *global* strategy were selected. We used Dijkstra’s algorithm to compute the shortest path from the starting position when the monkey switched to *global* strategy to the ending position when the monkey first reached a pellet. The trajectory with the fewest turns was determined by sorting all possible paths between the starting and the ending position.

### Eye Movement Analysis

We labelled monkeys’ fixation targets based on the distance between the eye position and the relevant game objects: Pac-Man, ghosts, pellets, and energizer. When the distances are within one tile (25 Pixel), we added the corresponding target to the label. There can be multiple fixation labels because these objects may be close to each other.

In Figure 3E, we selected strategy periods with more than ten consecutive steps and computed the fixation ratio by dividing the time that the monkeys spent looking at an object within each period by the period length. As pellets and energizers do not move but Pac-Man and ghosts do, we did not differentiate between fixations and smooth-pursuits when measuring where the monkeys looked at. We computed the average fixation ratio across the periods with the same strategies.

In the pupil dilation analyses in Figure 3F and Figure S6A, B, C, D, we z-scored the pupil sizes in each game round. Data points that were 3 standard deviations away from the mean were excluded. We aligned the pupil size to strategy transitions and calculated the mean and the standard error (Figure 3F, solid line and shades). As the control, we selected strategy transitions that go through the *vague* strategy and aligned the data to the center of the vague period to calculate the average pupil size and the standard error (Figure 3F, dashed line and shades). Figure S6A, B, C, D were plotted in a similar way but only specific strategy transitions.

### Compound Strategy Analysis

#### Planned Attack

We defined *planned attack* and *accidental consumption* trials according to the strategy labels after the energizer consumption: when at least eight out of ten time steps after the energizer consumption are labeled as the *approach* strategy, the trial is defined as *planned attack*; otherwise this trial is defined as *accidental consumption*. There were 493 (Monkey O) and 463 (Monkey P) *planned attack* trials and 1970 (Monkey O) and 1295 (Monkey P) *accidental consumption* trials. These trials are aligned to the time of energizer consumption in Figure 5A.

In Figure 5B, the average of Pac-Man-energizer distance, energizer-ghost distance, and Pac-Man-ghost distance is computed at the beginning of the *planned attack* and *accidental consumption* trials. The beginning of each trial is defined as the position where Pac-Man started to take the direct shortest route towards the energizer. The average fixation ratios in Figure 5C are computed from the beginning of each *planned attack* or *accidental consumption* till when the energizer is eaten.

In some of the *accidental consumption* trials (Monkey O: 590/30.3%, Monkey P: 478/36.9%), Pac-Man caught a ghost although the monkeys didn’t pursue the ghosts immediately after the energizer consumption. In contrast, all *planned attack* trials resulted in Pac-Man catching the ghosts successfully. These trials are aligned to the ghosts consumption in Figure 5D.

#### Suicide

We defined *suicide* and *failure* trials based on the strategy labels in the last ten steps before the Pac-Man’s death: a trial is defined as *suicide* when all 10 steps are labelled as *approach* and as *failure* when all steps are labelled as *evade*.

In Figure 6A, the distance between Pac-Man and the closest pellet and the distance between Pac-Man reset location and the ghost are computed at the time point when the monkeys switched to the *approach* (*suicide*) or *evade* (*failure*) strategy. Also, in Figure 6B, the average distance between the Pac-Man and two ghosts is computed in the same condition. The average fixation ratios in Figure 6C are computed from that time point until the Pac-Man’s death. In Figure 6D, the relative pupil sizes are aligned to the Pac-Man’s death.

**Figure S1.**
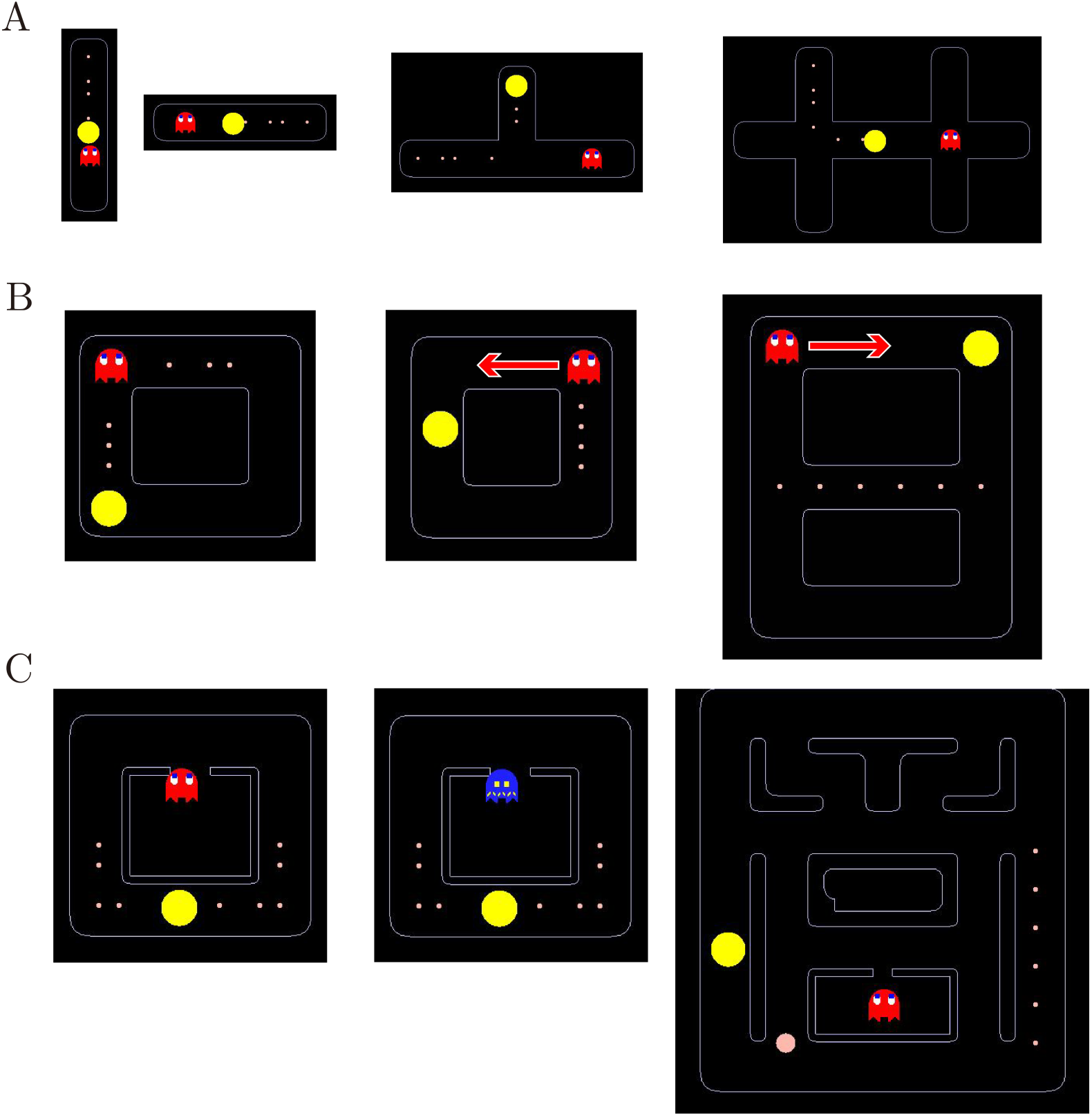
Training procedure. **A**. Stage one training mazes. From left to right: 1. Vertical maze. Pac-Man started from the middle position, with several pellets in one direction and a static ghost in the other. The monkeys learned to move the joystick upward and downward. 2. Horizontal maze. The monkeys learned to move the joystick toward left and right. 3. T-maze. Pac-Man started from the vertical arm, and the monkeys learned to move out of it by turning left or right. Pellets were placed in one arm and a static ghost in the other. 4. H-maze. Pac-Man started from the middle of the maze. There were pellets placed on the way leading to one of the three arms, and a static ghost was placed at the crossroad on the opposite side. **B**. Stage two training mazes. From left to right: 1. Square maze with a static ghost. Pac-Man started from one of the four corners, and pellets were placed in two adjacent sides with a static ghost placed at the corner connecting the two. 2. Square maze with a moving ghost. Pac-Man started from the middle of one of the four sides, and pellets were placed on the opposite side. A ghost moved from one end of the pellet side and stopped at the other end. 3. 8-shaped maze with a moving ghost. Pac-Man stated from one of the four corners. The pellets were placed in the middle tunnel. A ghost started from a corner and moved toward the pellets. **C**. Stage three training mazes. From left to right: 1. Square maze with Blinky. Pac-Man started from the middle of the bottom side with pellets placed on both sides. Blinky in normal mode started from its home. 2. Square maze with a ghost in a permanent scared mode. The scared ghost started from its home. Once caught by Pac-Man, the ghost went back to its home. 3. Maze with an energizer and Blinky. An energizer was randomly placed in the maze. Once the energizer was eaten, the ghost would be turned into the scared mode. The scared mode lasted until the ghost was eaten by Pac-Man. Once the ghost was eaten, it returned to its home immediately and came out again in the normal mode. After monkeys were able to perform the task, we limited the scared mode to 14 seconds.

**Figure S2.**
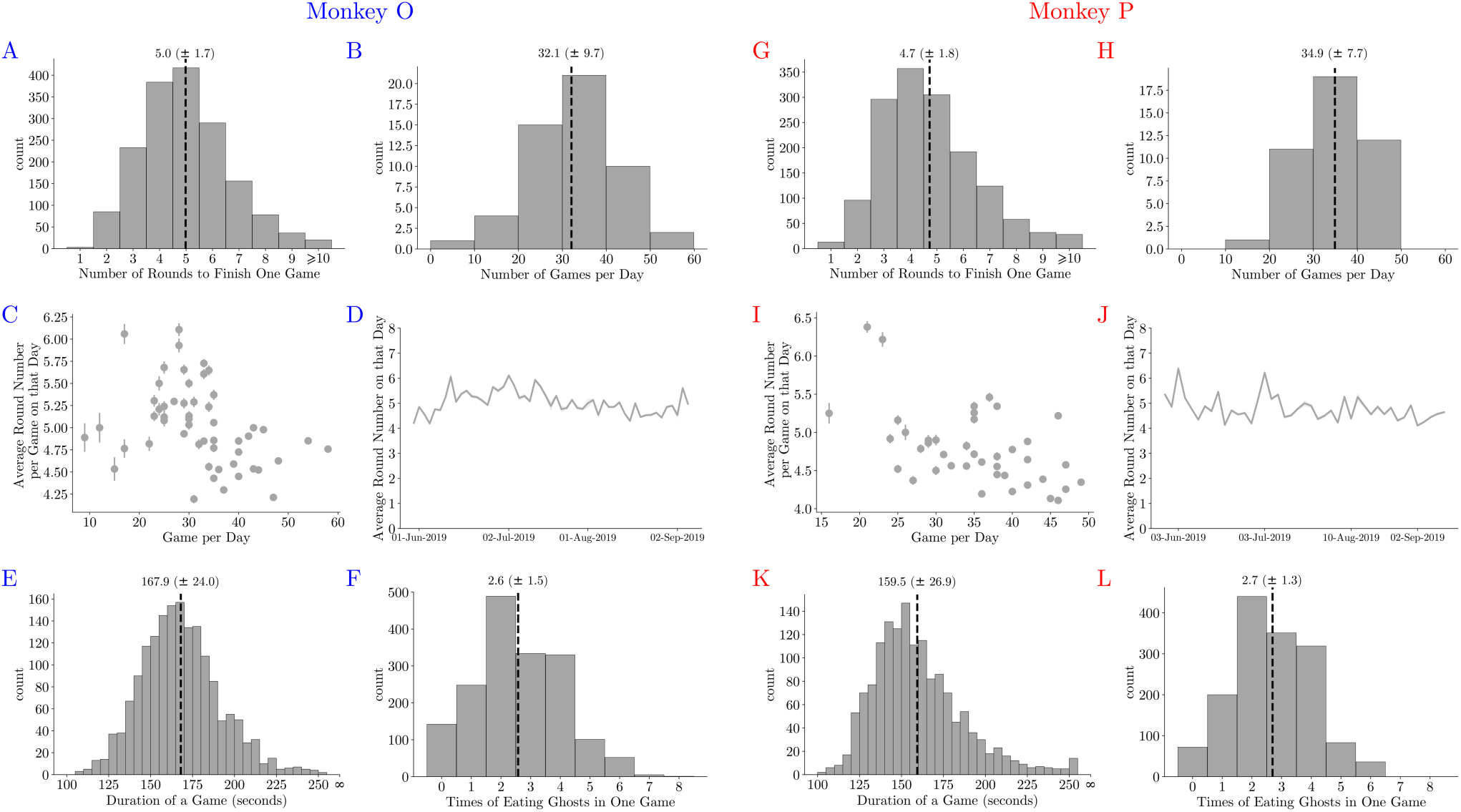
Basic game statistics of Monkey O (left) and Monkey P (right). **(A. G.)** The number of rounds to clear all pellets in each game. **(B. H.)** The number of games accomplished on each day. **(C. I.)** The average number of rounds to clear a maze plotted against the number of games in a session. Fewer numbers of rounds allowed the monkeys to complete more games in a session. **(D. J.)** The average number of rounds during the training. **(E. K.)** The time needed to clear a maze. **(F. L.)** The number of the ghosts that Pac-Man ate when completing a game.

**Figure S3.**
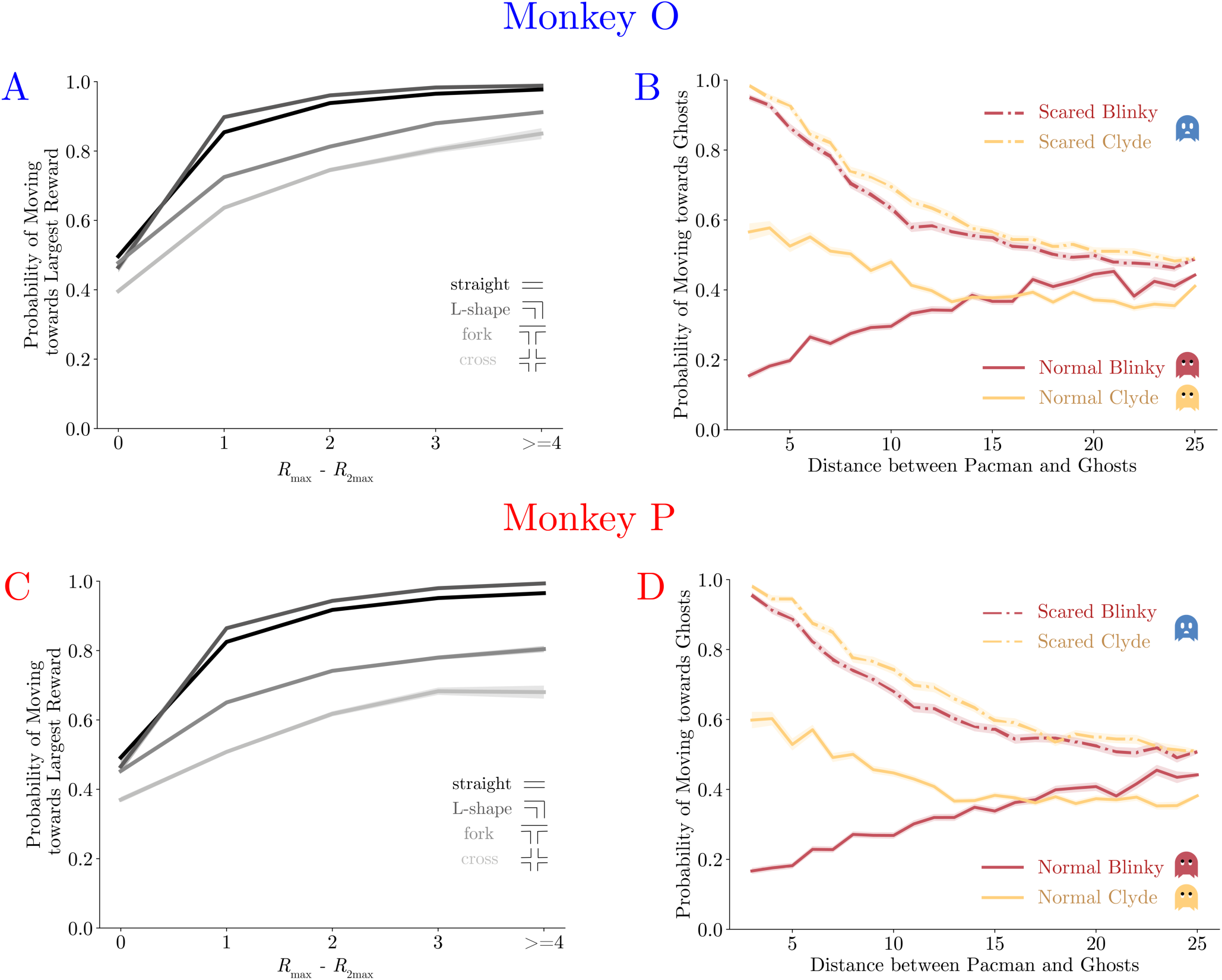
The performance of Monkey O (left) and Monkey P (right). (**A. C.)** Probability of monkeys moving towards the largest reward. The monkeys were more likely to move toward the direction with more local rewards. The abscissa is the reward difference between the most and the second most rewarding direction. The colors indicate the path types in which different numbers of moving directions are possible. (**B. D.)** Probability of monkeys moving towards ghosts. The monkeys escaped from the normal ghosts and chased the scared ghosts. The abscissa is the distance between Pac-Man and the ghosts. The colors and the line types indicate the ghosts and their modes.

**Figure S4.**
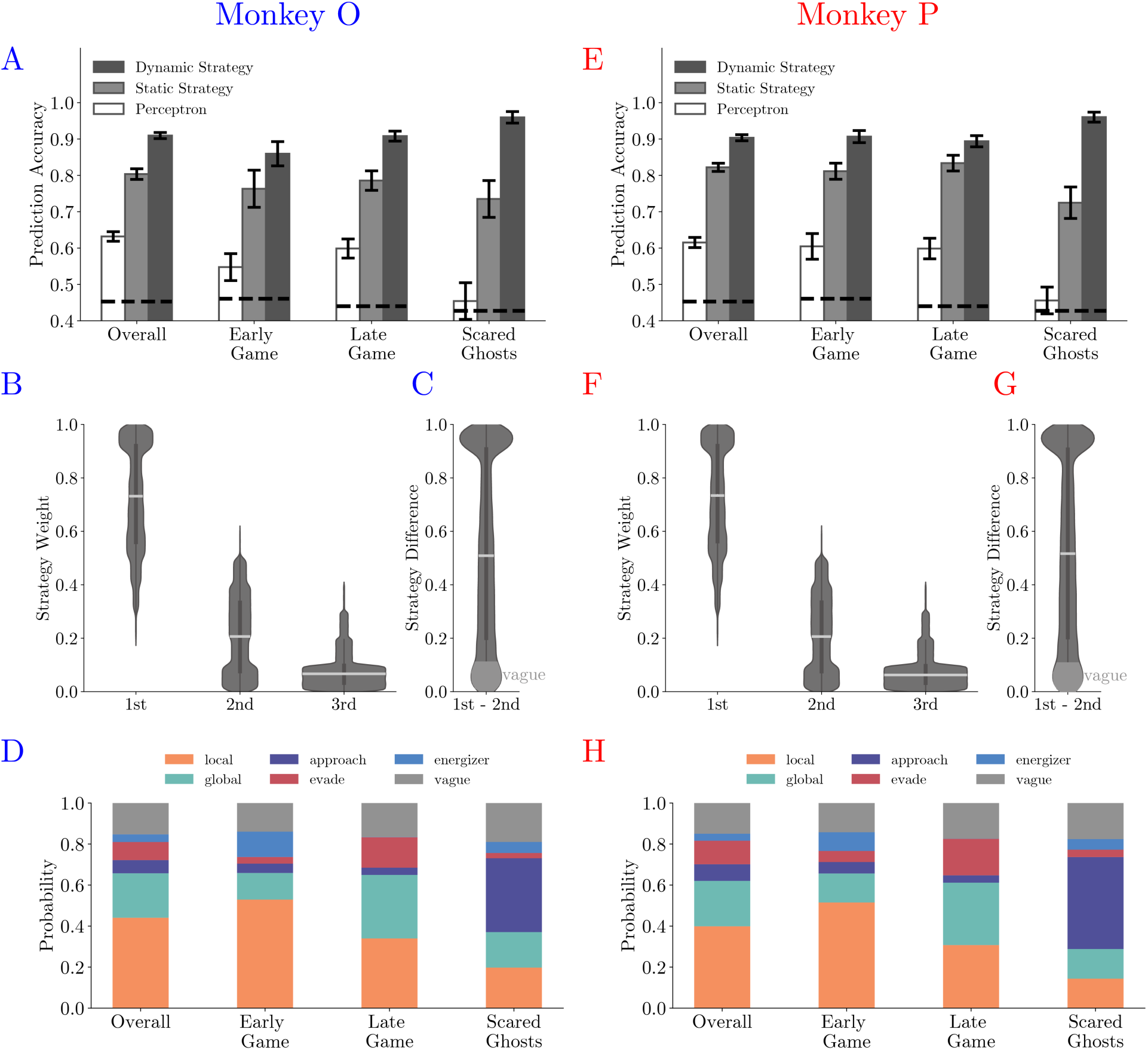
Fitting behavior with strategy labels for Monkey O (left) and P (right). **(A. E**.) Comparison of prediction accuracy across three models in four game contexts. See **Table S5** and **Table S6** for details. **(B. F.)** The histograms of the three dominating strategies’ weights. **B**. The most dominating strategy’s weights (0.735±0.210) were much larger than the secondary strategy (0.201±0.156) and tertiary strategy (0.052±0.079). **F**. The most dominating strategy’s weights (0.737±0.208), the secondary strategy (0.201±0.156), and tertiary strategy (0.050±0.078). **(C. G.)** The histogram of the three dominating strategies’ weights. In over 85% of the time, the weight difference was larger than 0.1, and more than a quarter of the time the difference was over 0.9. **(D. H.)** The ratios of labeled dominating strategies across four game contexts. In the early game, the *local* strategy was the dominating strategy. In comparison, in the late game, both the *local* and the *global* strategies had large weights. The weight of the *approach* strategy was largest when the ghosts were in the scared mode.

**Figure S5.**
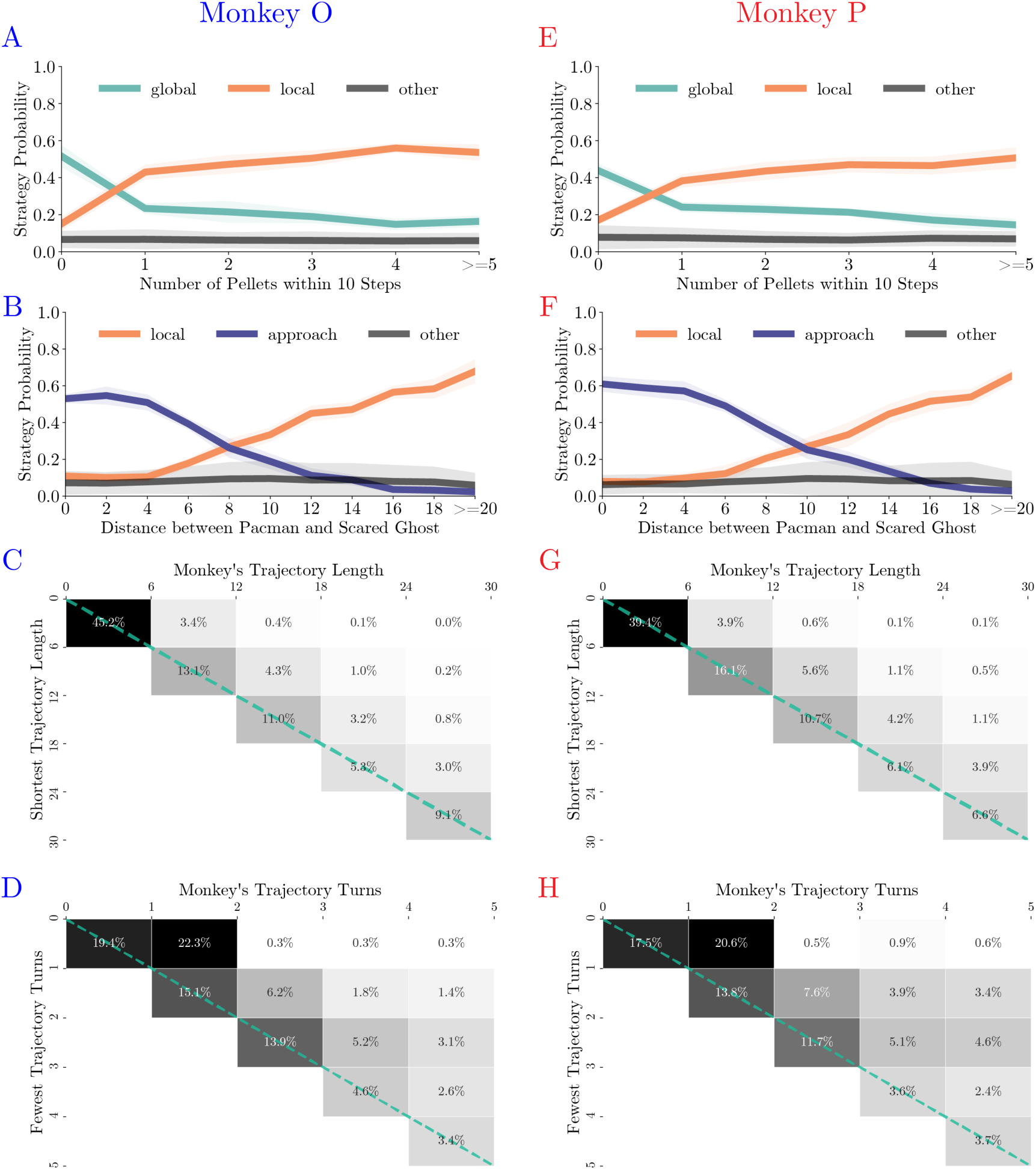
Monkey’s behavior under different strategies for Monkey O (left) and P (right). **(A. E.)** The probabilities of the monkeys adopting the *local* or *global* strategy correlate with the number of *local* pellets. **(B. F.)** The probabilities of the monkeys adopting the *local* or *approach* strategy correlate with the distance between Pac-Man and the ghosts. **(C. G.)** When adopting the *global* strategy to reach a far-away patch of pellets, the monkeys’ actual trajectory length was close to the shortest. The column denotes the actual length, and the row denotes the optimal number. The percentages of the cases with the corresponding actual lengths are presented in each cell. High percentages in the diagonal cells indicate close to optimal behavior. **(D. H.)** When adopting the *global* strategy to reach a far-away patch of pellets, the monkeys’ number of turns was close to the fewest possible turns. The column denotes the actual turns, and the row denotes the optimal number. The percentages of the cases with the corresponding optimal numbers are presented in each cell. High percentages in the diagonal cells indicate close to optimal behavior.

**Figure S6.**
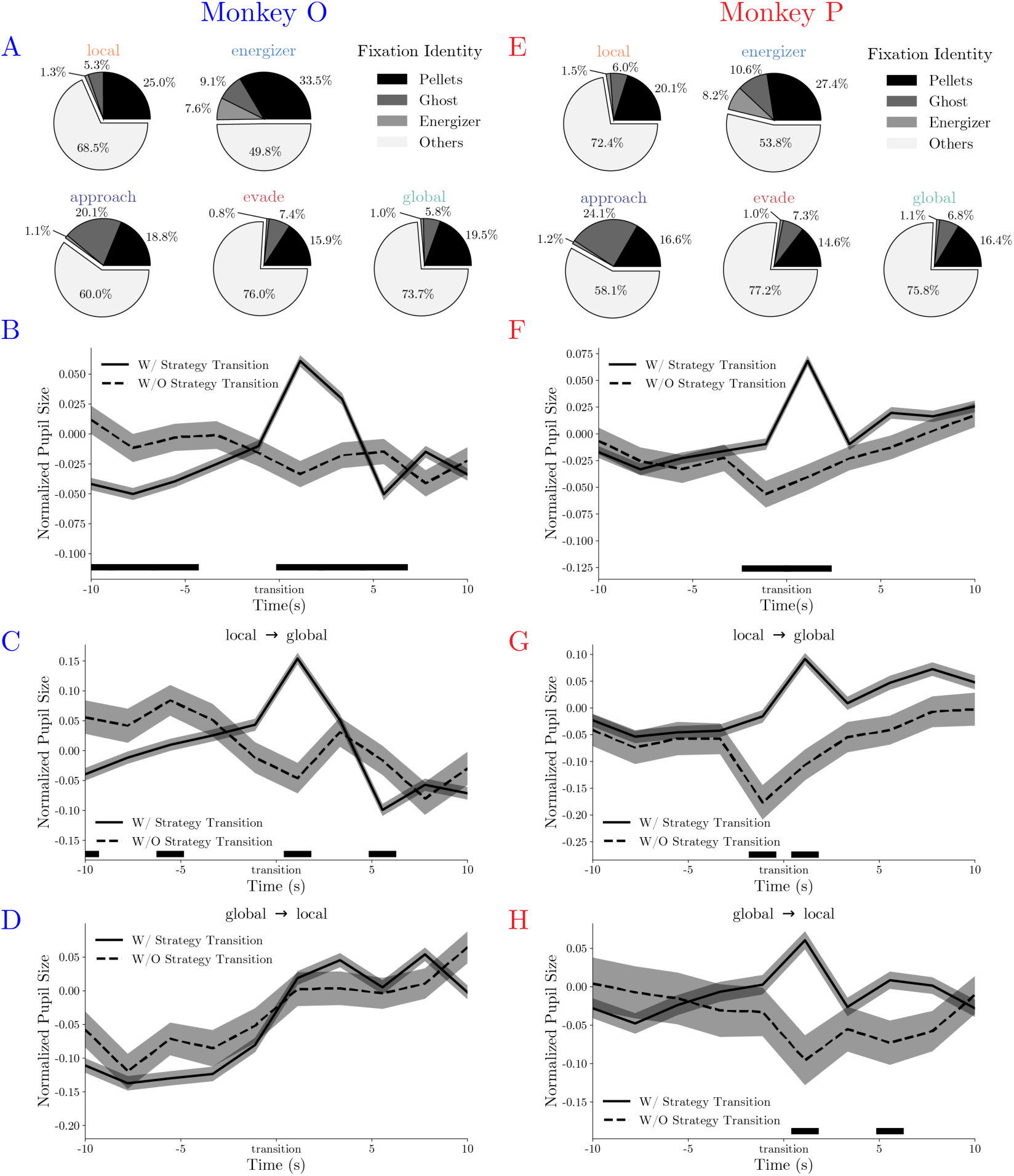
Monkey’s eye movement patterns under different strategies for Monkey O (left) and P (right). **(A. E.)** Average fixation ratios of ghosts, energizers, and pellets when the monkeys used different strategies. **(B. F.)** The monkeys’ pupil diameter increases around the strategy transition (solid line). Such increase was absent if the strategy transition went through the *vague* strategy (dashed line). Black bars at the bottom denote p < 0.01, two-sample t-test. **(C. G.)** The Monkeys pupil diameter increase was evident in transitions from *local* to *global*. The shades are standard errors across trials. Black bars at the bottom denote p < 0.01, two-sample t-test. **(D. H.)** The Monkeys pupil diameter increase was evident in transitions from *global* to *local* in monkey P. It was not obvious in monkey O, as the pupil size increased in general. The shades are standard errors across trials. Black bars at the bottom denote p < 0.01, two-sample t-test.

**Figure S7.**
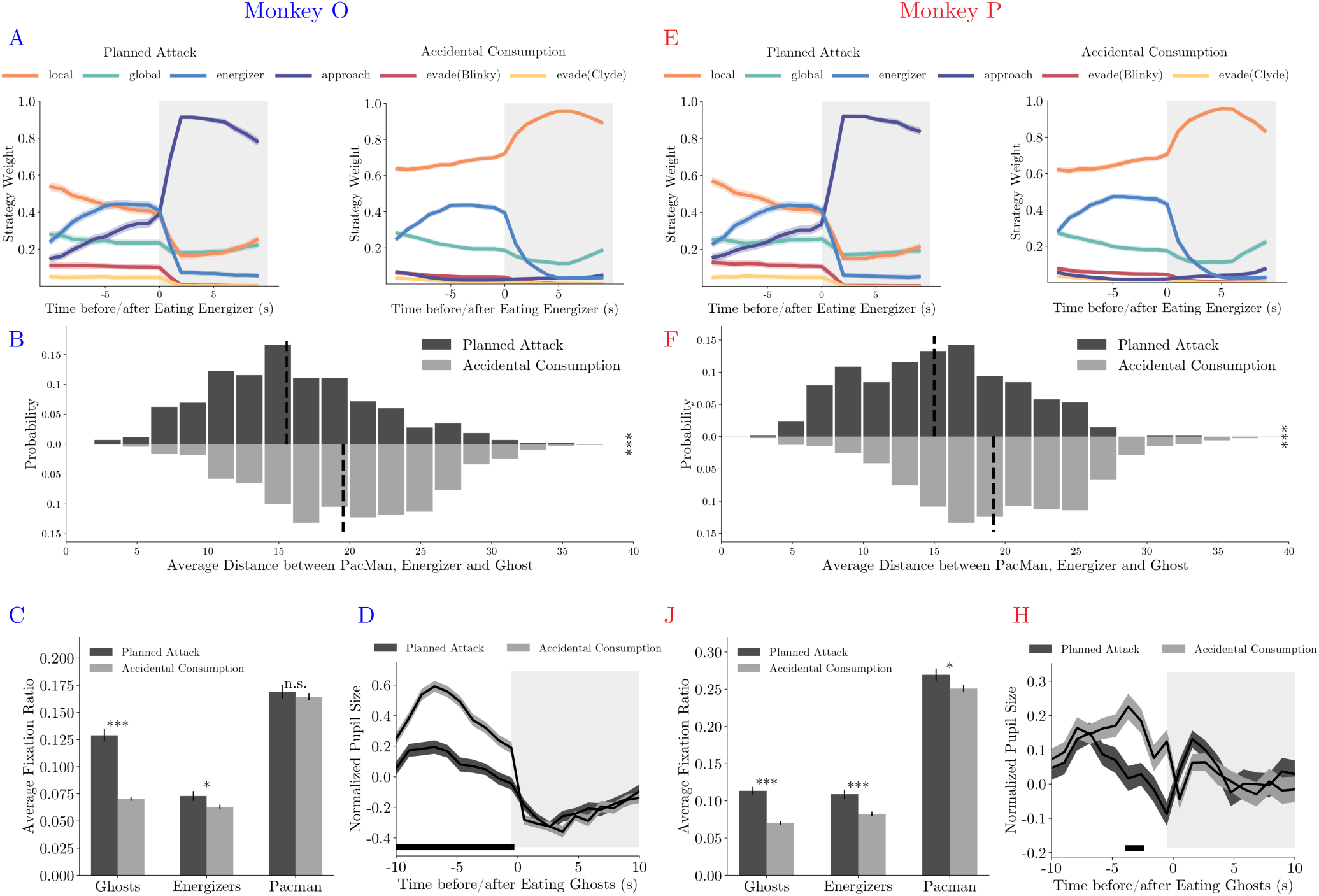
*Planned attacks* in Monkey O (upper) and P (lower). **(A. E.)** Average strategy weight dynamics in *planned attacks* (left) and *accidental consumptions* (right). **(B. F.)** The average distance between the Pac-Man, the energizer, and the ghosts in *planned attacks* and *accidental consumptions*. *** denotes p<0.001, two-sample t-test. **(C. J.)** Ratios of fixations on the ghosts, the energizer and the Pac-Man. *** denotes p<0.001 and ** denotes p<0.01, two-sample t-test. **(D. H.)** The pupil size aligned to the ghost consumption. The black bar near the abscissa denotes data points where the two traces are significantly different (p < 0.01, two-sample t-test).

**Figure S8.**
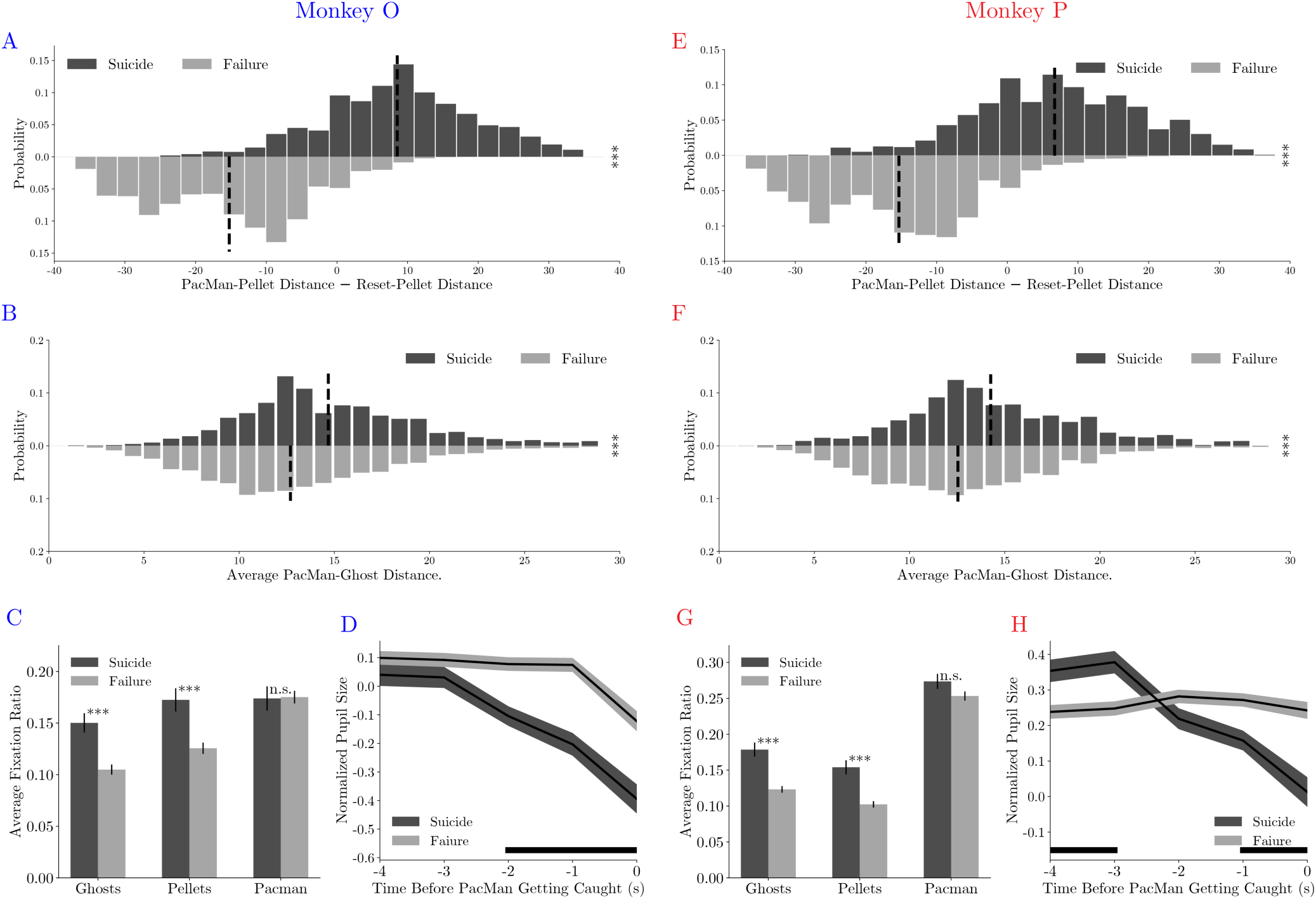
*Suicides* in Monkey O (left) and P (right). (**A. E.)** The distance difference between Pac-Man and closest pellet before and after the death are smaller in *suicides* than in *failed evasions*. *** denotes p<0.001, two-sample t-test. **(B. F.)** The average distance between Pac-Man and the ghosts was greater in *suicides* than in *failed evasions*. *** denotes p<0.001, two-sample t-test. **(C. G.)** The monkeys fixated more frequently on the ghosts and the pellets in *suicides* than in *failed evasions*. *** denotes p<0.001, two-sample t-test. **(D. H.)** The monkeys’ pupil size decreased before the Pac-Man’s death in *suicides*. The black bars near the abscissa denote data points where the two traces are significantly different (p < 0.01, two-sample t-test).

**Table S1.** Four path types in the maze.

| Path Type | Selection Criteria |
| --- | --- |
| straight | Contains two opposite moving directions |
| L-shape | Contains two orthogonal moving directions |
| fork | Contains three moving directions |
| cross | Contains four moving directions |

**Table S2.** Awarded and penalized utilities for each game element in the model.

| Reward($r_p$) | Reward($r_e$) | Reward($r_f$) (1-5) | Reward($g$) | Penalty($g$) |
| --- | --- | --- | --- | --- |
| 2 | 4 | 3, 5, 8, 12, 17 | 8 | -8 |

**Table S3.**
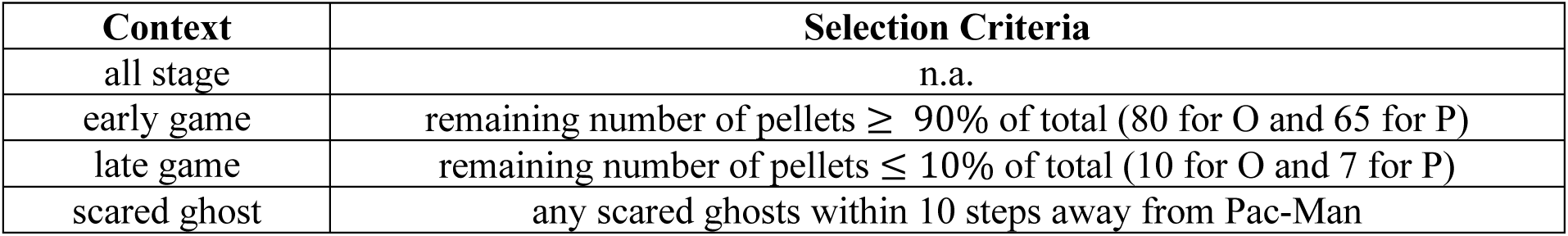
Special game contexts and corresponding selection criteria.

| Context | Selection Criteria |
| --- | --- |
| all stage | n.a. |
| early game | remaining number of pellets ≥ 90% of total (80 for O and 65 for P) |
| late game | remaining number of pellets ≤ 10% of total (10 for O and 7 for P) |
| scared ghost | any scared ghosts within 10 steps away from Pac-Man |

**Table S4.** Comparison of prediction accuracy (± SE) across three models in four game contexts (Table S3) for two monkeys.

| Strategy<br>Context | Dynamic | Static | Perceptron |
| --- | --- | --- | --- |
| Overall | 0.907 ± 0.006 | 0.813 ± 0.010 | 0.624 ± 0.010 |
| Early Game | 0.889 ± 0.017 | 0.792 ± 0.024 | 0.582 ± 0.026 |
| Late Game | 0.902 ± 0.010 | 0.808 ± 0.018 | 0.599 ± 0.019 |
| Scared Ghosts | 0.960 ± 0.010 | 0.729 ± 0.033 | 0.455 ± 0.030 |

**Table S5.** Comparison of prediction accuracy (± SE) across three models in four game contexts (Table S3) for monkey O.

| Strategy<br>Context | Dynamic | Static | Perceptron |
| --- | --- | --- | --- |
| Overall | 0.910±0.008 | 0.804±0.015 | 0.632±0.013 |
| Early Game | 0.860±0.033 | 0.763±0.051 | 0.548±0.037 |
| Late Game | 0.910±0.014 | 0.786±0.027 | 0.598±0.026 |
| Scared Ghosts | 0.960±0.016 | 0.735±0.051 | 0.545±0.050 |

**Table S6.** Comparison of prediction accuracy (± SE) across three models in four game contexts (Table S3) for monkey P.

| Strategy<br>Context | Dynamic | Static | Perceptron |
| --- | --- | --- | --- |
| Overall | 0.904±0.008 | 0.822±0.011 | 0.615±0.014 |
| Early Game | 0.907±0.017 | 0.811±0.022 | 0.605±0.035 |
| Late Game | 0.894±0.015 | 0.834±0.022 | 0.599±0.028 |
| Scared Ghosts | 0.960±0.01 | 0.725±0.043 | 0.456±0.037 |

**Movie S1-S10 are uploaded at YouTube: https://youtube.com/playlist?list=PL_iWMTlE8pjv6mnKVjOaaO02hMKHm2AAj**

**Movie S1-S3**. Example game trials. Monkey O’s moving trajectory, actual and predicted actions, and labeled strategies were plot in these example game trials. In these trials, the monkey can solve the task with one attempt.

**Movie S4-S5**. Example game trials. Monkey P’s moving trajectory, actual and predicted actions, and labeled strategies were plot in this example game segment. In these trials, the monkey can solve the task with one attempt.

**Movie S6**. Example game segment. Monkey’s moving trajectory, actual and predicted actions, and labeled strategies were plot in this example game segment. Monkey’s real-time saccade position was plot as a moving white dot. In this example, the monkey started with the *local* strategy and grazed pellets. With the ghosts getting close, it initiated the *energizer* strategy and went for a nearby energizer. Once eating the energizer, the monkey switched to the *approach* strategy to hunt the scared ghosts. Afterward, the monkey resumed the *local* strategy and then used the *global* strategy to navigate towards another patch when the local rewards were depleted.

**Movie S7**. *Planned attack* game segment. Monkey’s moving trajectory, actual and predicted actions, and labeled strategies were plot in this *planned attack* game segment. Monkey’s real-time saccade position was plot as a moving white dot. In this trial, the monkey immediately switched to the *approach* strategy after eating an energizer.

**Movie S8**. *Accidental consumption* game segment. Monkey’s moving trajectory, actual and predicted actions, and labeled strategies were plot in this *accidental consumption* game segment. Monkey’s real-time saccade position was plot as a moving white dot. In this trial, the monkey treated an energizer just as a more rewarding pellet and continued collecting pellets with the *local* strategy after eating the energizer.

**Movie S9**. *Suicide* game segment. Monkey’s moving trajectory, actual and predicted actions, and labeled strategies were plot in this *suicide* game segment. Monkey’s real-time saccade position was plot as a moving white dot. In this trial, the monkey moved Pac-Man towards a normal ghost to be eaten on purpose. The local pellets were scarce, and the remaining pellets were far away. Thus, the death of Pac-Man resets the game and returns Pac-Man and the ghosts to their starting positions, making *suicide* a more advantageous compound strategy.

**Movie S10**. *Failed evasion* game segment. Monkey’s moving trajectory, actual and predicted actions, and labeled strategies were plot in this *failed evasion* game segment. Monkey’s real-time saccade position was plot as a moving white dot. In this trial, the monkeys were adopting the *evade* strategy but failed to escape from the ghosts.

